# Multi-omic integration with human DRG proteomics highlights TNFα signalling as a relevant sexually dimorphic pathway

**DOI:** 10.1101/2024.12.06.626968

**Authors:** Allison M Barry, Julia R Sondermann, Joseph B Lesnak, Feng Xian, Úrzula Franco-Enzástiga, Jayden A O’Brien, David Gomez-Varela, Morgan K Schackmuth, Stephanie Shiers, Theodore J Price, Manuela Schmidt

**Affiliations:** Department of Neuroscience and Center for Advanced Pain Studies, University of Texas at Dallas, Richardson, TX 75080, USA; Systems Biology of Pain, Division of Pharmacology & Toxicology, Department of Pharmaceutical Sciences, University of Vienna, Austria

## Abstract

The peripheral nervous system (PNS) plays a critical role in pathological conditions, including chronic pain disorders, that manifest differently in men and women. To investigate this sexual dimorphism at the molecular level, we integrated quantitative proteomic profiling of human dorsal root ganglia (hDRG) and peripheral nerve tissue into the expanding omics framework of the PNS. Using data-independent acquisition (DIA) mass spectrometry, we characterized a comprehensive proteomic profile, validating tissue-specific differences between the hDRG and peripheral nerve. Through multi-omic analyses and *in vitro* functional assays, we identified sex-specific molecular differences, with TNFα signalling emerging as a key sexually dimorphic pathway with higher prominence in males. Genetic evidence from genome-wide association studies (GWAS) further supports the functional relevance of TNFα signalling in the periphery, while clinical trial data and meta-analyses indicate a sex-dependent response to TNFα inhibitors. Collectively, these findings underscore a functionally sexual dimorphism in the PNS, with direct implications for sensory and pain-related clinical translation.

## Introduction

Neuroimmune- and pain- related condition prevalence, as well as the corresponding treatment efficacy, can differ across sex and gender [28,41]. Understanding sexual dimorphism at a molecular level is thus a fundamental clinical issue. Autoimmune conditions, for example, are highly biased toward women (upwards of 80%), and infections can elicit differing immune responses. Women also have heightened sensitivity to pain in an experimental context, show higher rates of chronic pain, and are the dominant gender in many pain-centric disorders, from migraine to complex regional pain syndrome (CRPS) [8,15,41].

Pre-clinical work in rodents shows a similar sexual dimorphic trend at the immune and nervous system level. Decades of work suggest complex, cross-species mechanism(s) underlying these clinical presentations. Hormones such as oestrogen, prolactin, and testosterone have been implicated in a range of pain and neuro-immune conditions, while evidence for central and peripheral sexually dimorphic mechanisms have been shown [32,45,51,52,62].

Primary afferents in the peripheral nervous system (PNS) project from the skin and viscera to the dorsal horn of the spinal cord, with cell bodies in the dorsal root ganglia (DRG, in humans - hDRG). The DRG contain sensory neuron cell bodies, as well as a diverse set of non-neuronal cells. These sensory neurons respond to inflammation by detecting immune mediators and then releasing neuropeptides in complex neuro-immune circuits [33,58]. At a molecular level, transcriptional profiles of these cell types have recently been published for human sensory neurons [9,55], as well as non-neuronal cells [9,34] with some differences in gene expression reported across male and female donors. In bulk tissue, recent molecular profiling of the hDRG has highlighted differentially accessible chromatin regions (DARs), as well as sex-specific changes in neuropathic pain states at the transcriptome level [25,45]. To date, the corresponding proteome signature has been missing, as the few prior studies depend on shotgun (data dependent acquisition) proteomics [19,50], which is known to be biased towards high-abundant proteins [27].

Here, we generate a quantitative hDRG proteomic dataset across male and female donors using data-independent acquisition (DIA) mass spectrometry. This comprehensive dataset provides a human-centric reference for protein expression in a key structure of the peripheral nervous system, and complements previously published ATAC-seq and RNA-seq data. Through integration across omics from the same tissue, we show strong evidence for sexual dimorphism in the TNFα signalling pathway. Genome-wide association studies (GWAS) provide support that it is a functionally relevant pathway in the periphery, and evidence from clinical trials highlights a link to clinical sexual dimorphism in response to medication against these targets. The functional relevance of TNFα signalling in the hDRG was confirmed *in vitro*, and downstream changes in protein phosphorylation suggests a possible mechanism of action. Together, these findings speak to a sexually dimorphic pathway in the peripheral nervous system, which is of particular importance to sensory- and pain-related translation.

## Methods

### DRG tissue procurement

All human tissue procurement procedures were approved by the Institutional Review Board at the University of Texas at Dallas. Through collaboration with the Southwest Transplant Alliance, human lumbar DRGs (hDRGs, L1-L4) from organ donors were obtained within 4 hours of cross-clamp. Tissue for mass spectrometry was frozen in dry ice right immediately and stored in a −80°C freezer. All transport was done on large amounts of dry ice to protect tissue integrity. Age-matched male and female samples of mixed ethnicity were used for the current study. Donors for proteomic profiling were negative for pain, neuropathy, and illicit drug use (excluding marijuana), based on recorded patient history.

For Fluorescent Activated Cell Sorting (FACS) hDRGs (lumbar and thoracic) were recovered and stored in 10 mL of Hibernate A (BrainBits, HACA500) supplemented with 1% N-2 (Thermo Scientific, 17502048), 2% NeuroCult SM1 (Stemcell technologies, 05711), 1% penicillin/streptomycin (Thermo Fisher Scientific, 15070063), 1% Glutamax (Thermo Scientific, 35050061), 2mM Sodium Pyruvate (Gibco, 11360-070), and 0.1% Bovine Serum Albumin (Biopharm, 71-040) at 4⁰C until ready for processing (10-16 hrs, see below).

### DRG tissue preparation for mass spectrometry

DRGs were processed sequentially, with experimenters blinded to a randomized order (corresponding to donor column, Table 1). 12-24 hours prior to dissection, tissue was transferred to -20°C. For each hDRG, tissue was thawed on ice for ∼15 min prior to dissection (on ice) to remove the surrounding connective tissue. In each ganglia, a region of nerve tissue extending from the core ganglia was identified. This was presumed to contain fewer/no neuronal soma and was transected crudely with a scalpel for separate processing (as the “nerve root” region).

**Table 1.** Donor details.

| donor | Sex | Age | DRG | Pain | HTN | Ethnicity | COD |
| --- | --- | --- | --- | --- | --- | --- | --- |
| MS-5 | F | 32 | L2 | FALSE | FALSE | Black | Head Trauma/GSW |
| MS-3 | F | 41 | L3 | FALSE | TRUE | White | Head Trauma/Motorcycle Accident |
| MS-7 | F | 41 | L2 | FALSE | TRUE | White | CVA/Stroke |
| MS-2 | F | 55 | L3 | FALSE | TRUE | White | Anoxia/Cardiovascular |
| MS-1 | M | 33 | L3 | FALSE | TRUE | White | Head Trauma/MVA |
| MS-8 | M | 41 | L4 | FALSE | TRUE | Black | Head Trauma/GSW/Suicide |
| MS-4 | M | 43 | L3 | FALSE | FALSE | Hispanic | Head Trauma/MVA |
| MS-6 | M | 53 | L1 | FALSE | FALSE | Hispanic | Anoxia/Seizure |
| FACS-1 | M | 34 | 2xL3+T8 | FALSE | FALSE | Black | Anoxia |
| FACS-2 | M | 37 | L1-3, L5 | TRUE | FALSE | White | Head Trauma/non-MVA |
| FACS-3 | M | 31 | L3 | FALSE | FALSE | Asian | Anoxia |
| FACS-4 | F | 29 | L1+L3 | FALSE | TRUE | White | Sepsis |
| FACS-5 | F | 41 | L3-L5 | NA | NA | NA | NA |

*MS: mass spectrometry; FACS: fluorescent activated cell sorting; HTN: hypertension, COD: cause of* *death; GSW: gunshot wound; MVA: motor vehicle accident; CVA: cerebrovascular accident*

### Protein extraction and SP3-assisted digestion

Due to the size of the ganglia, each ganglia was cut into up to 4 pieces and subsequently, each piece was cut in smaller pieces to increase surface area for lysis. All small pieces of one big cut were transferred to a 2ml LoBind Protein Eppendorf tube (Eppendorf, Hamburg, Germany) prefilled with lysis buffer (2% SDS, 100mM Tris, 5% glycerol, 10mM DTT and 1x protease inhibitor cocktail) and sonicated for 15 min (cycles of 30 sec ON and 30 sec OFF, 4°C, low frequency) in a Bioruptor Pico (Diagenode, Seraing, Belgium). Samples were subsequently incubated for 15 min at 70°C, 1500 rpm in a ThermoMixer C (Eppendorf) followed by a centrifugation for 5 min with 10000 × g at room temperature (RT) to pellet cell debris. The supernatant was mixed with 5x sample volume 100% acetone (pre-chilled at -20°C) and incubated for 2.15-2.45 h to precipitate the proteins.

Precipitates were pelleted by centrifugation for 30 min at 14000 × g, RT. Pellets were washed with ice-cold 80% EtOH and centrifugated again with above mentioned parameters. The EtOH was removed and pellets air-dried for around 20 min at RT. Resolubilization in lysis buffer was facilitated by incubation for 10 min at 70°C, 1500 rpm. Total protein concentration was determined at 280 nm with a NanoPhotometer N60 (Implen, Munich, Germany). Subsequently, samples were flash-frozen and stored at -20°C until further usage.

Protein clean-up and digest was then performed as described in Xian et al., 2022. This is based on the single-pot, solid-phase-enhanced sample preparation (SP3) method from Hughes et al. (Hughes et al., 2019), and is also detailed in the corresponding protocol at protocol.io [5]. Remaining protein was used for a targeted phosphorylation array (below).

### Tandem mass spectrometry

Nanoflow reversed-phase liquid chromatography (Nano-RPLC) was performed on NanoElute2 systems (Bruker Daltonik, Bremen, Germany). This was coupled with timsTOF HT (Bruker Daltonik, Bremen, Germany) via CaptiveSpray ion source. Mobile phase A consisted of 100% water, 0.1% formic acid and mobile phase B of 100% acetonitrile, 0.1% formic acid. 500 ng of peptides were loaded onto a C18 trap column (1 mm x 5 mm, ThermoFisher) and further separated over a 90 min gradient on an AuroraTM ULTIMATE column (25 cm × 75 µm) packed with 1.6 µm C18 particles (IonOpticks, Fitzroy, Australia). The flow rate was set to 250 nL/min, except for the last 7 minutes, where the flow rate was accelerated to 400 nL/min. The mobile phase B was linearly increased from 2 to 20% in the first 60 minutes, followed by another linear increase to 35% within 22 minutes and a steep increase to 85% in 0.5 min. Then, a flow rate switch to 400 nL/min was achieved in 0.5 min and was maintained for 7 minutes to the end of the gradient to elute all hydrophobic peptides.

The samples were analyzed in data-independent acquisition (DIA) mode coupled with parallel accumulation serial fragmentation (PASEF). Precursors with m/z between 350 and 1200 were defined in 13 cycles (either 2 or 3 quadrupole switches per cycle) containing 34 ion mobility steps within the ion mobility range of 0.65 – 1.35 (1/k0) with fixed isolation window of 25 Th in each step. The acquisition time of each DIA-PASEF scan was set to 100 ms, which led to a total cycle time of around 1.48 sec. The collision energy was ramped linearly from 65 eV at 1/k0 = 1.6 to 20 eV at 1/k0 = 0.6.

### Spectral deconvolution with DIA-NN

DIA-NN (version 1.8.1) was used to process raw spectra in library-free mode via command line on the Vienna Scientific Cluster [16,17]. A predicted library search was performed against the human proteome (UP000005640) with match-between-runs (MBR) enabled. Separate MBRs were performed for quality control samples, and ganglia + nerve root, as well as ganglia and nerve root independently when used for differential expression testing. Gene Group outputs are also referred to as “proteins” throughout. See data availability statement for access.

### Basic Gene Lists

Curated lists of pain genes have been previously published [36,40,64]. Ion channels were matched from Alexander et al., (2023), and GO-term related gene lists were extracted through R using biomaRt [20]. Drug-targets were extracted from opentargets.com for relevant conditions [42]. Receptor types were derived from a previously published interactomics resource [58], and were compared to published bulk RNA-seq from the DRG [45].

### Gene Set Enrichment Analysis (GSEA)

Gene Set Enrichment Analysis (GSEA) for neuronal subtype enrichment was performed against gene lists derived from Zheng et al., [65] for mouse subpopulations as described previously [7]. In brief, RNA-seq count data from GSE131230 - which had been processed via STAR alignment and HTSeq on the same genome build (see Zheng et al., for full methods) - was corrected for library size and transformed via rlog in R using DESeq2 [38,66]. This was then filtered to match their published report. Genes with an average rlog above the 95% quantile cut-off per subpopulation were curated into a ‘gene set’ for enrichment. Mouse gene names were converted using the biomaRt package via the ensembl mart with ‘getLDS()’ [20].

For human subpopulations as described in Taveres-Ferreira et al., [55], marker genes were accessed directly from the supplemental tables and reformatted in R for processing. In brief, these genes were selected using the ‘findMarkers()’ function in Seurat after Visium-based spatial sequencing.

These custom gene sets were then compiled for a GSEA analysis using the clusterProfiler package [61]. Minimum gene set size was set to 25 (no maximum size threshold), and run for 10000 permutations.

The average expression across ganglia samples was calculated per gene group after filtering for 80% completeness. Where the first term per gene group overlapped (eg. TRPV1 vs TRPV1,TRPV2,TRPA1), the first term in the group was used, and the row with the higher abundance was taken. GSEA was then performed on the ranked mean expression.

To examine pathway differences between male and female samples, GSEA was performed on ranked LFC output with a significance threshold of FDR < 0.05; minimum gene set size was set to 25. Gene sets were extracted using the ‘msigdbr’ package in R [18]. Hallmark pathways were extracted as ‘category = “H”’. Biological pathways (BP) and molecular function (MF) from gene ontology (GO) lists were extracted as ‘category = “C5”, subcategory = “BP”’ and ‘ category = “C5”, subcategory = “MF”’ respectively.

For the proteomics data, LFC were calculated using limma (see: hypothesis testing). For GSEA on RNA-seq, DESeq2 was used to calculate LFC, as described in Farah et al., (2024), using previously published bulk RNA-seq data (n = 31M, 19F donors) [45]. For ATAC-seq, LFC were derived from pseudobulk analysis of all clusters in a spatial-seq dataset in autosomes, described by Franco-Enzástiga et al., (2024). In brief, ATAC-seq gene scores were calculated using ArchR. The gene score predicts the expression of genes based on the chromatin accessibility of regulatory elements in the vicinity of a gene and its corresponding gene body. In ArchR, the function ‘GeneScoresMatrix()’ was used to store gene scores. To sum together gene scores of all the cells from the same sample for the pseudobulk analysis, the ‘getMarkerFeatures()’ function in ArchR was used, and the ‘groupBy = “Sample”’ was specified. Data were compared across sex (n = 5M, 3F donors).

### Overrepresentation Analysis

Overrepresentation analysis was performed using the ‘enrichGÒ function from ClusterProfiler [61]. Differentially expressed proteins (DEPs, abs(LFC) > 1, FDR < 0.05) were compared to a background of Gene Groups used for the initial limma comparison using default settings.

### Supervised PCA (SPCA)

Supervised PCA (SPCA) [3] was performed as previously described [7]. Differentially expressed genes (DEGs) from spatial RNA-seq and differentially expression regions (DARs) from bulk hDRG ATAC-seq datasets exploring sexual dimorphism were extracted as lists [25,55]. DARs were filtered to remove X and Y chromosome regions. DEGs from Taveres-Ferreira et al., (2022) were extracted from supplemental table 2, ‘B-Overall_neurons_DE_genes’ looking at differential expression between male and female barcodes within neurons.

Proteomic data from the ganglia (merged by replicate) was subset by either DEGs or DARs and subject to a PCA (via ‘prcomp’ in R). Eigengenes and eigen vectors were then extracted from the first principal component (’ PC1’).

### Hypothesis Testing

Hypothesis testing was performed within ganglia samples using a moderated T-Test via limma [46], modelling for sex, n = 4M and 4F donors. Here, log2 transformed data were first filtered for the primary occurrence of each Gene Group (ie. protein) and filtering out proteins present in less than 80% of samples. For example, where Gene Group A = “TRPV1” and B = “TRPV1,TRPV2,TRPA1”, Gene Group A would be used. P values were corrected with Benjamini-Hochberg, and a significance threshold was set to FDR < 0.05, LFC >1.

### Multi-study Factor Analysis

Strong correlations between ATAC-seq regions, RNA-seq transcript abundance, and protein abundance were not evident. Even so, matching sexually dimorphic pathways are described across datasets. Here, we employed a Multi-study factor analysis [57] to examine latent variables in the context of sex to look for the underlying structure contributing to this overlap.

Following the principles for factor analyses on small sample sizes [59] we limited our search to a small number of factors (here, 3) and used sexually dimorphic DARs. This allows us to look exclusively at chromatin regions implicated in female/male differences: this biases the analysis towards factors involved in sexual dimorphism and provides a clear link from ATAC-RNA-protein. The within-dataset factor numbers were determined by scree plots. A minimum Kaiser– Meyer–Olkin (KMO) threshold was set to 0.5 for each dataset, and only sex-related factors were considered from the output (ie., a confirmatory model, opposed to an exploratory factor analysis). Due to the arbitrary nature of factor loading signs, within-dataset factor loadings were constrained such that they were positively correlated to males if sex appeared relevant.

Bulk RNA-seq from human DRG [45] were extracted as quantile-normalized transcripts per million (qnTPM) for neuronal-enriched donor samples (n = 50). Both “pain” and “non-pain” donors were considered here to examine sexual dimorphism across states. Proteomic samples from the “ganglia” and “nerve root” regions of the DRG were used, merged across technical replicates by mean. Here, we acknowledge the limitation that this is a low sample number, and that samples are paired (16 samples from 8 donors). We limited this effect by analyzing only factors shared across two datasets, specifically in the context of sex, and not within the proteomic dataset in isolation.

Genes were matched to bulk DARs from Franco-Enzástiga et al., (2024) and filtered to remove genes with missing values across both datasets, resulting in complete data matrices. Data were then transposed for Multi-study factor analysis (RNA = [50 x 402], proteomics = [16 x 402]). Starting values were extracted using ‘start_msfà, constraint = ‘block_lower2’, capped at 10000 iterations. ‘ecm_msfà was then run with default parameters.

Factor scores were estimated using the Weighted Least Squares (ie. the Bartlett Method; equation 1), where X∼ is the centred/scaled data matrix for each dataset, *i*. Lambda are the factor loadings shared across datasets and Psi is a diagonal matrix equal to the specific variances.

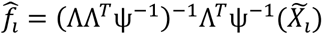

The interaction of ‘Sex’ and ‘Factor’ was tested by an anova. Posthoc testing was then performed per factor (eg. Male.Factor1 - Female.Factor1) using ‘glht(anova, linfct = mcp(Interaction))’ in R. Factors with a q < 0.1 were considered significant. Shared gene loadings were extracted from the Multi-Study Factor Analysis output as Phi for visualization.

GSEA analyses were performed as above, with the following modification: minimum gene set size set to 5 (due to the total number of genes available, 402). In line with above, the significance threshold was set to FDR < 0.05. Protein-protein interaction networks were plotting through https://livedataoxford.shinyapps.io/drg-directory/, [64] which uses an API with STRING DB and overlays sensory neuron and pain-relevant datasets through R Shiny.

### Fluorescent Activated Cell Sorting (FACS) preparation

The DRGs were trimmed of excess connective tissue, fat, and nerve roots to collect the bulb containing neuronal cell bodies. The bulb of each DRG was cut into 3mm sections and placed in 5mL of pre-warmed digestion enzyme containing 2 mg/mL of Stemxyme I (Worthington Biochemical, LS004106), 10ng/mL of recombinant human β-NGF (R&D Systems, 256-GF), and 0.1 mg/mL of DNAse I (Worthington Biochemical, LS002139) in HBSS without calcium and magnesium (Thermo Scientific, 14170-112).

The tubes were placed in a 37° C shaking water bath and triturated every hour until the DRG sections were dissolved (3-4hrs). Samples were filtered through a 70µm mesh strainer and centrifuged at 350g for 5 minutes at room temperature (all subsequent centrifuge steps follow the same parameters). The supernatant was removed, and the pellet was resuspended in a red blood cell lysis buffer (Biolegend, 420301) and incubated at room temperature for 5 minutes. Samples were then centrifuged, the supernatant was removed, and the pellet was resuspended in 0.5% bovine serum albumin in 1X Phosphate Buffered Saline (PBS). To remove myelin from the dissociation, cells were incubated with myelin removal beads (Miltenyi Biotec, 130-096-433) for 15 minutes at room temperature. Cells were washed with 1mL of 0.5% bovine serum albumin in PBS, spun down, resuspended in 0.5% bovine serum albumin in PBS, and passed through a LS column (Miltenyi, 130-042-401) on a MidiMACS separator (130-042-301) according to manufacturer’s protocol. Samples were then spun down and resuspended in PBS and proceeded to cell staining.

Cells were stained with a fixable live/dead stain (Biolegend, 423107) for 10 minutes at room temperature, protected from light. Cells were washed with 1mL of flow cytometry staining buffer (Invitrogen, 00-4222-26), spun down, and resuspended in flow buffer. Cells were incubated with an Fc receptor blocker (TruStain FcX, Biolegend, 422302) for 10 minutes at room temperature, protected from light. Cells were then incubated with CD45, CD11b, and CD3 antibodies for 30 minutes on ice, protected from light (See Supplemental Table 10 for antibodies used in flow cytometry experiments). Cells were washed with 1mL of flow cytometry staining buffer, spun down, and resuspended in flow buffer and kept on ice until processing. Fluorescently activated cell sorting (FACS) was used to isolate Live, CD45+, CD11b+, and CD3-cells on a BD FACSAria Fusion (Gating Strategy in Supplemental Figure 9A). Cells were collected, spun down, and resuspended in pre-warmed RPMI media (Gibco, 11875-093) containing 10% HyClone™ Fetal Bovine Serum (Thermo Fisher Scientific, SH3008803IR) and 1% penicillin/streptomycin. Cells were plated at 20k cells per well in a 96 well plate and allowed to acclimate overnight in an incubator (37⁰C, 5% CO2). All antibodies are listed in Supplemental Table 10.

### LPS Stimulation of hDRG immune cells

Cells were stimulated with lipopolysaccharide (LPS) (10ng/mL, Sigma-Aldrich, L6529) or its vehicle (RPMI media) for 16 hours as this dose has previously shown to drive increased secretion of TNFα from human myeloid cells *in vivo* [1,11]. In one condition cells were co-stimulated with Brefeldin A (1:1000, Biolegend, 420601). In cultures without Brefeldin A, the cell culture supernatant was collected and frozen at -80°C till processing. TNFα levels in the media supernatant were measured with a Legendplex kit targeting TNFα (Biolegend, 741187), per the manufacture’s recommended protocol. The average of two technical replicates were used to calculate TNFα concentrations. In cells co-cultured with Brefeldin A, cells were collected and fixed with a Cyto-Fast fixation and permeabilization kit (Biolegend, 426803) for 20 minutes at room temperature. Cells were washed twice with the Cyto-Fast perm wash solution and then incubated with a TNFα antibody for 30 minutes on ice. Cells were washed with 1mL of flow cytometry staining buffer, spun down, resuspended in flow buffer, and kept on ice until data acquisition on a BD LSRFortessa. The percentage of cells expressing TNFα were calculated for each condition using FlowJo (Version v10.10) (Gating Strategy in Supplemental Figure 9B).

### TNFα Stimulation

hDRG immune cells were stimulated with TNFα (10ng/mL, R&D Systems, 210-TA) or its vehicle (1X PBS) for 30 minutes. This dose and timepoint has been shown to increase p38 levels in human cultures of myeloid cells and fibroblasts [30,54]. Cells were then collected and fixed with 4% paraformaldehyde (Electron Microscopy Sciences, 15710) in PBS for 15 minutes at room temperature. Cells were then washed 2x with flow cytometry staining buffer and permeabilized with pre-chilled, 100% methanol (Fisher Scientific, A456-212), for 30 minutes on ice. Cells were then washed 2x with flow cytometry staining buffer and incubated with p38 and p65 antibodies in flow buffer for 30 minutes on ice. Cells were washed with 1mL of flow cytometry staining buffer, spun down, resuspended in flow buffer, and kept on ice until processing. The percentage of cells expressing p38 and p65 along with the median fluorescent intensity of each signal was calculated for each condition (Gating Strategy in Supplemental Figure 9B). The median fluorescent intensity values were normalized to an unstained control.

### Targeted Phosphorylation Array

Phosphorylation levels were measured in remaining hDRG protein samples using the Proteome Profiler Human Phospho-Kinase Array Kit (R&D Systems, CAT# ARY003C). The commercial protocol was followed, with the following changes: each membrane was washed independently using 7 ml Wash Buffer in 60x15 mm culture dishes, and membranes were rinsed 1x with wash buffer after antibody incubation prior to 3x10 min washes. A ChemiDoc MP Imaging Platform (Bio Rad) was used to detect signals. Four samples per condition were processed for membrane “A”, while 3F + 4M were processed for membrane “B”.

Briefly, the protocol is as follows: membranes were blocked with “Array Buffer 1” for 1 hr at RT prior to an overnight incubation at 4°C with 250 ug protein per membrane (ie. 500 ug per membrane pair, A&B), diluted to 1 ml in “Array Buffer 1”. Membranes were then washed (3x10 min), incubated with membrane-specific antibodies for 2 hr at RT, rinsed 1x prior to washing (3x10 min), and incubated with Streptavidin-HRP for 30 min at RT. After washing (3x10 min), membrane were incubated for 90 seconds with the “Chemi Reagent Mix”, blotted with Kimwipes and imaged on a ChemiDoc MP Imaging Platform for 240s. All membranes per round (2M/2F per round) were developed and imaged together.

Image quantification was performed in ImageJ. Regions of Interest (ROIs) of equal size were processed using the ROI Manager, and quantification was considered as 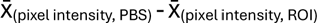 for each phosphorylation site, using the PBS negative control per membrane as the background.

### Graphics

All plots were generated in R with the following libraries unless otherwise described: ggplot2, ComplexHeatmap, cowplot, gridExtra, ggbiplot, ggrepel. Figures were compiled in Inkscape.

### Results

In this study, we present the most comprehensive proteomic dataset of the human dorsal root ganglia (hDRG) and the nerve root (Fig 1A), with previous datasets using shotgun proteomics [19,50]. Using data independent acquisition coupled with parallel accumulation serial fragmentation (DIA-PASEF) we identify ∼ 12500 Gene Groups (proteins) per sample, with similar levels between the ganglia and root (Fig 1B-C, see Data Availability). Donor details are available in Table 1. We confirmed data quality using control samples throughout the MS runs for both LC-MS and SP3 replicates (SFig 1A), and do not see overt variance by donor or age (SFig 1B). As expected, samples show the highest correlation within sample replicates (SFig 1C). Even so, samples do not cluster obviously by tissue type or donor (Fig 1D-E) or by sex/replicate (SFig 1D), suggesting a generalized proteome signature across all samples. These technical replicates were merged (by mean), with 32 samples (16x2 replicates) averaged to 8 ganglia + 8 nerve samples for downstream analyses (from 4M + 4F donors).

**Figure 1.**
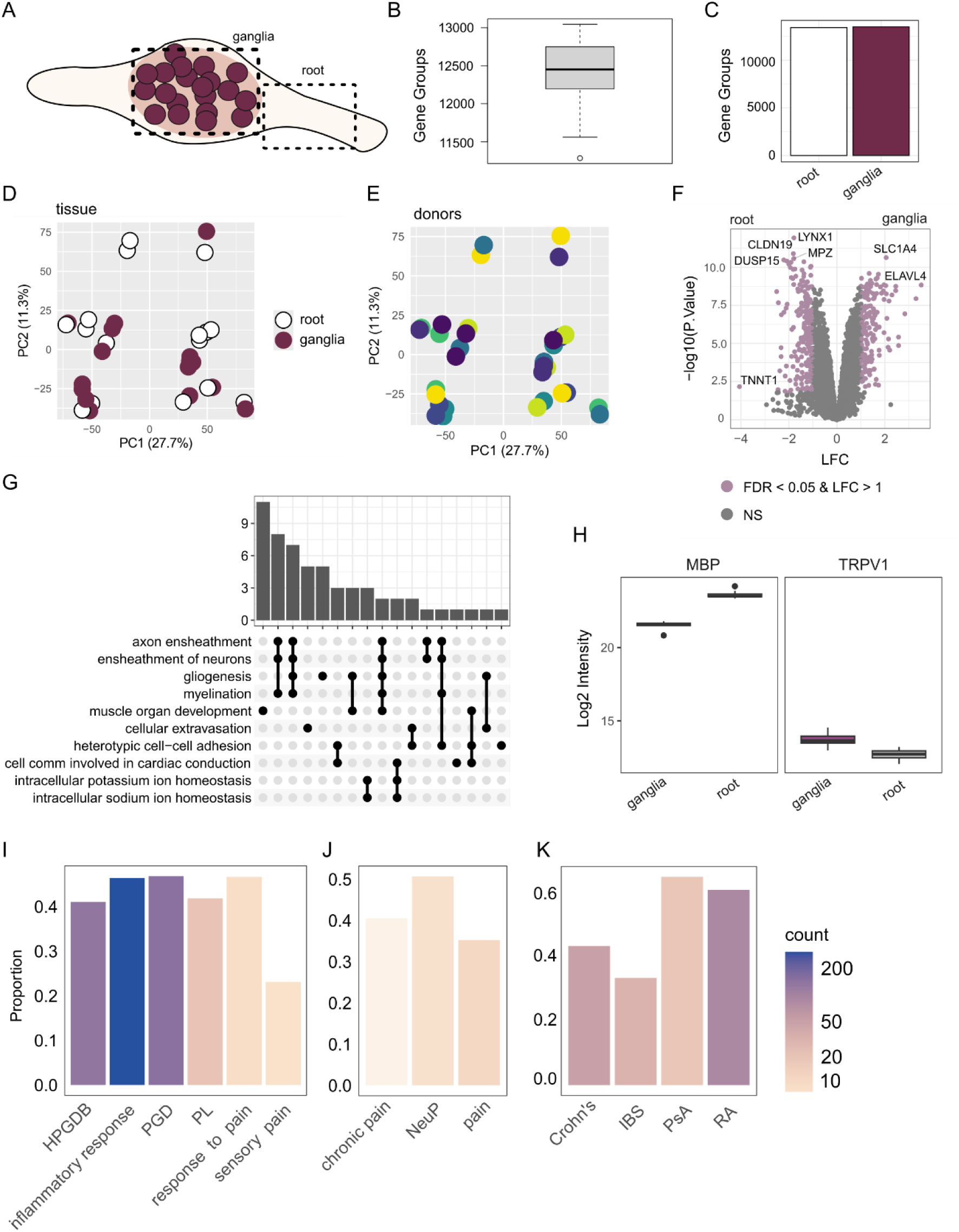
Quantitative proteomics of the hDRG and nerve root. A. Schematic of the ganglia and nerve root (NR, root). B. Gene Groups (protein counts) per sample. C. Gene Group count per tissue. D-E: PCA for tissue (D) and donor (E). F. Volcano plot of differentially expressed proteins (DEPs, ganglia vs nerve root; positive LFC for ganglia), coloured by abs(LFC) and FDR < 0.05. NS: not significant. G. Upset plot showing GO term over enrichment analysis for differentially expressed proteins upregulated in the nerve root. H. Example DEPs for myelin basic protein (MBP) and TRPV1 (adjusted p < 0.01 each). I-K. Overlap of detected Gene Groups with relevant gene sets: related to pain (I), pain drugs (J), peripheral autoimmune conditions (K). HPGDB: Human Pain Genetics Database [40], PGD: Pain Gene Database [36], PL: DoloRisk priority group Pain List [56]. NeuP: neuropathic pain. IBS: Irritable Bowel Syndrome, PsA: Psoriatic Arthritis, RA: Rheumatoid Arthritis. Related to SFig 1. n = 8 donors (4 M + 4F) for ganglia + nerve, with technical replicates (x2 per sample) merged for downstream analyses.

When comparing the nerve root to the ganglia, we see large differences in protein (aka “Gene Group”; see methods) levels. These differentially expressed proteins (DEPs) show a bias in myelin-associated terms upregulated in the root and soma-related terms in the ganglia (Fig 1F-H). A full list of DEPs is provided in Supplemental Table 1, with GO enrichment in Supplemental Tables 2-3. This is expected, given our knowledge of these tissues, and gives confidence to this dataset to probe other questions. With our current dataset quality and coverage, we see this data as a reference for the hDRG proteome, with extensive coverage of relevant curated gene lists for pain targets (Fig 1I), pain-related drug targets (Fig 1J), and peripheral autoimmune targets (Fig 1K).

Given the importance of membrane proteins in neurological and neuro-immune conditions in the PNS, as well as the previous difficulties in detecting membrane proteins using mass spectrometry, we specifically investigated ion channel and receptor abundance in our dataset (Fig 2) [50]. Here, we detect a number of TRP-, SCN-, and KCN-ion channels (Fig. 2A), among others, as well as a small number of GPCRs (Fig 2B). These candidates are expressed throughout our dynamic range of protein abundance (Fig 2C), with a number of key proteins detected almost exclusively in the ganglia (eg. P2RX3, SCN10A, Fig 2D). Additionally, when we compare these data to a curated list of proteins for sensory neurons and myelin, we see a clear separation of tissue types, again highlighting the strength of this dataset (Fig 2D).

**Figure 2.**
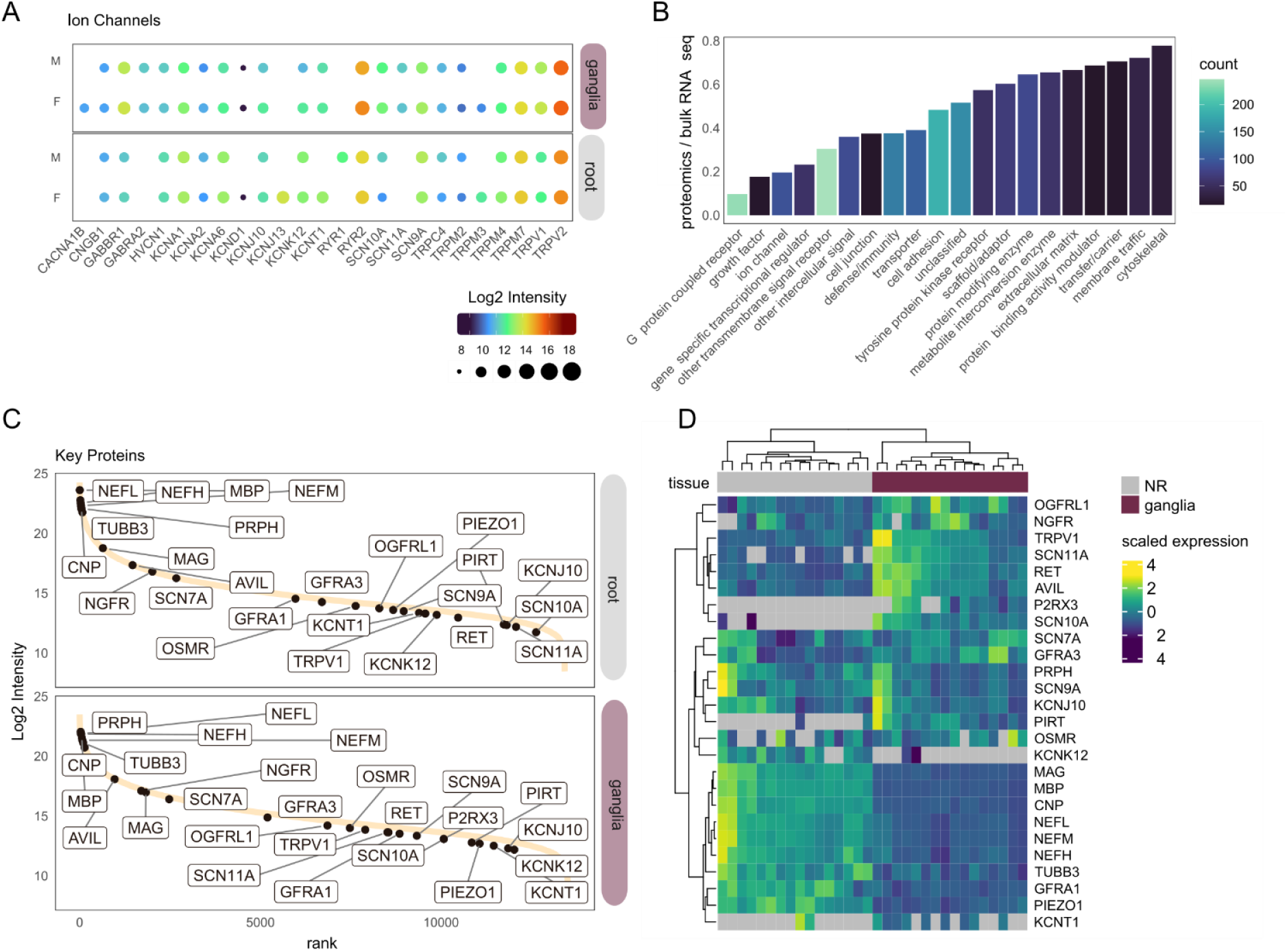
Ion channel and G-protein coupled receptor (GPCR) expression across tissues. A. Average expression by ion channel across tissues. B. Proportion of receptor detection in the proteomics data vs bulk RNA-seq of the hDRG [45]. C. Dynamic range plot showing ranked intensity across tissues. Key proteins (gene groups) for sensory neurons and myelin are highlighted. D. Heatmap across all samples (including each technical replicate) highlighting the dataset completeness for key proteins in C. n = 8 donors (4 M + 4F), with samples for ganglia + nerve; each sample has a technical replicate.

### The Ganglia and Nerve Root

Because of the differences between the ganglia and nerve root (Fig 1F-H), we independently investigated the proteome of each tissue type. In the ganglia (Fig 3A), we again do not see obvious clustering by PCA over cause of death and Ethnicity (Fig 3B-C). The samples cluster by replicate (SFig 2A) and show consistent protein counts (SFig 2B). In line with previous reports for quantitative proteomics, the data also does not correlate strongly to bulk RNA-seq of the same tissue [45], with an R^2^ = 0.15 (SFig 2C). When looking more precisely at receptor types, we see a highly variable correlation across receptor types, ranging from -0.2 to 0.8 (SFig 2D), likely reflecting a combination of biological and technical differences in receptor type proteomics, due in part to regulatory dynamics.

**Figure 3.**
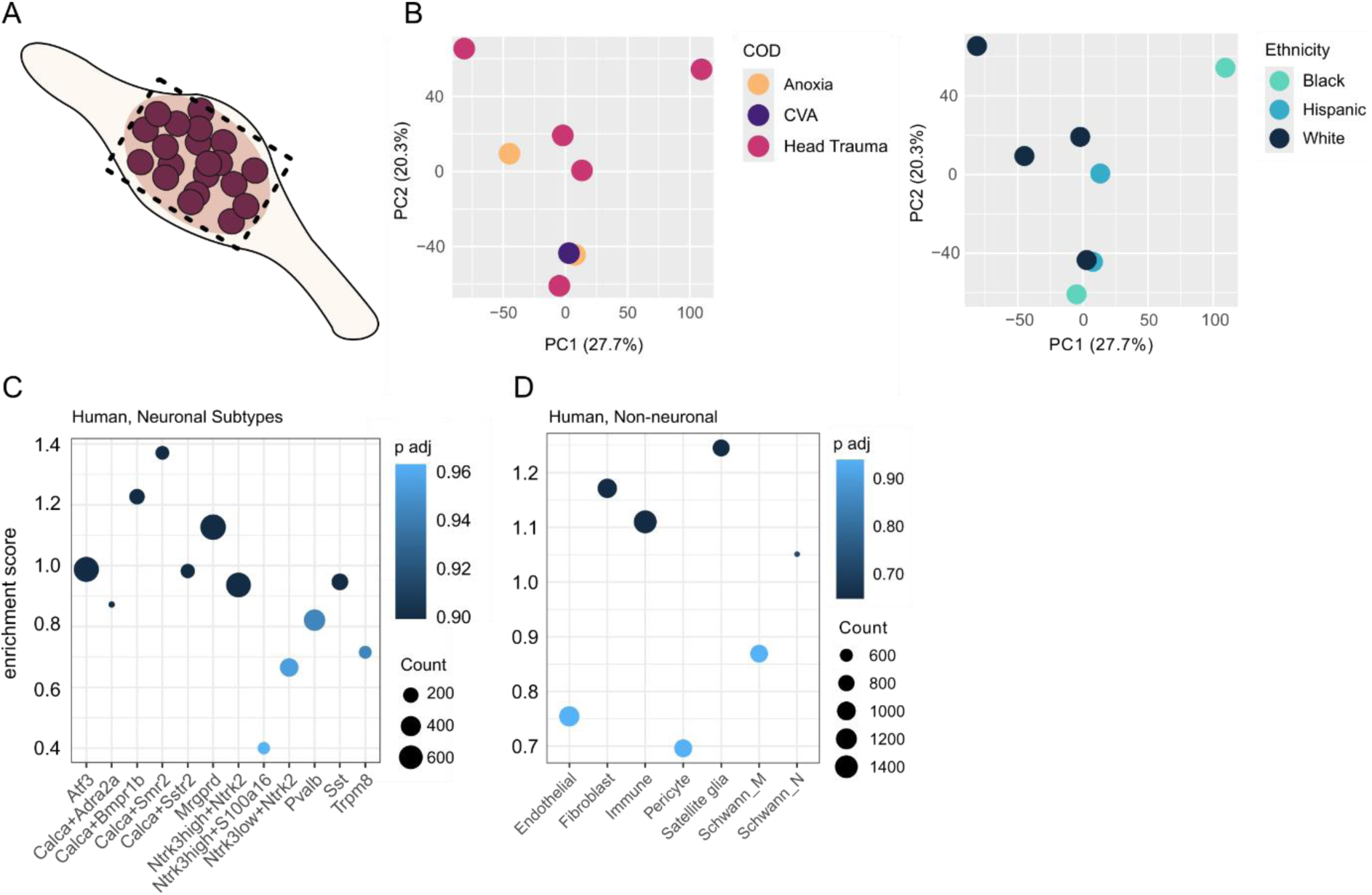
DIA-PASEF proteomics of the hDRG has markers across cell types. A. Schematic. B. PCA, coloured by cause of death (COD, left) and Ethnicity (right). C. Gene Set Enrichment Analysis (GSEA) against human neuronal subtype markers. D. GSEA against non-neuronal cell type markers. Related to SFig. 2-3. n = 4M + 4F donor DRGs.

Next, we examined the enrichment profiles of neuronal and non-neuronal cells using custom genesets derived from snRNA-seq [9] (Fig 3D, see: methods). Here, we do not see a specific significant enrichment for any cell type, but instead can detect marker genes across populations. This trend is mirrored when using gene sets derived from mouse datasets [65] (SFig 3A-B), and for neurons using a secondary, visium-based, hDRG dataset [55] (SFig 3C).

We then set out to investigate male/female differences in the hDRG (Fig 4, SFig 4). Ganglia samples do not cluster by sex (Fig 4A) and no differentially expressed proteins were detected between males and females (SFig 4A, Supplemental Table 4-5). Even so, gene set enrichment analysis (GSEA) highlights pathway differences (Fig 4B-C, SFig 4B-D). Notably, TNFα signalling is enriched in male donors (Fig 4B), while interferon (IFN) α response (Fig 4C) and Oxidative Phosphorylation (SFig 4D) are enriched in female samples.

**Figure 4.**
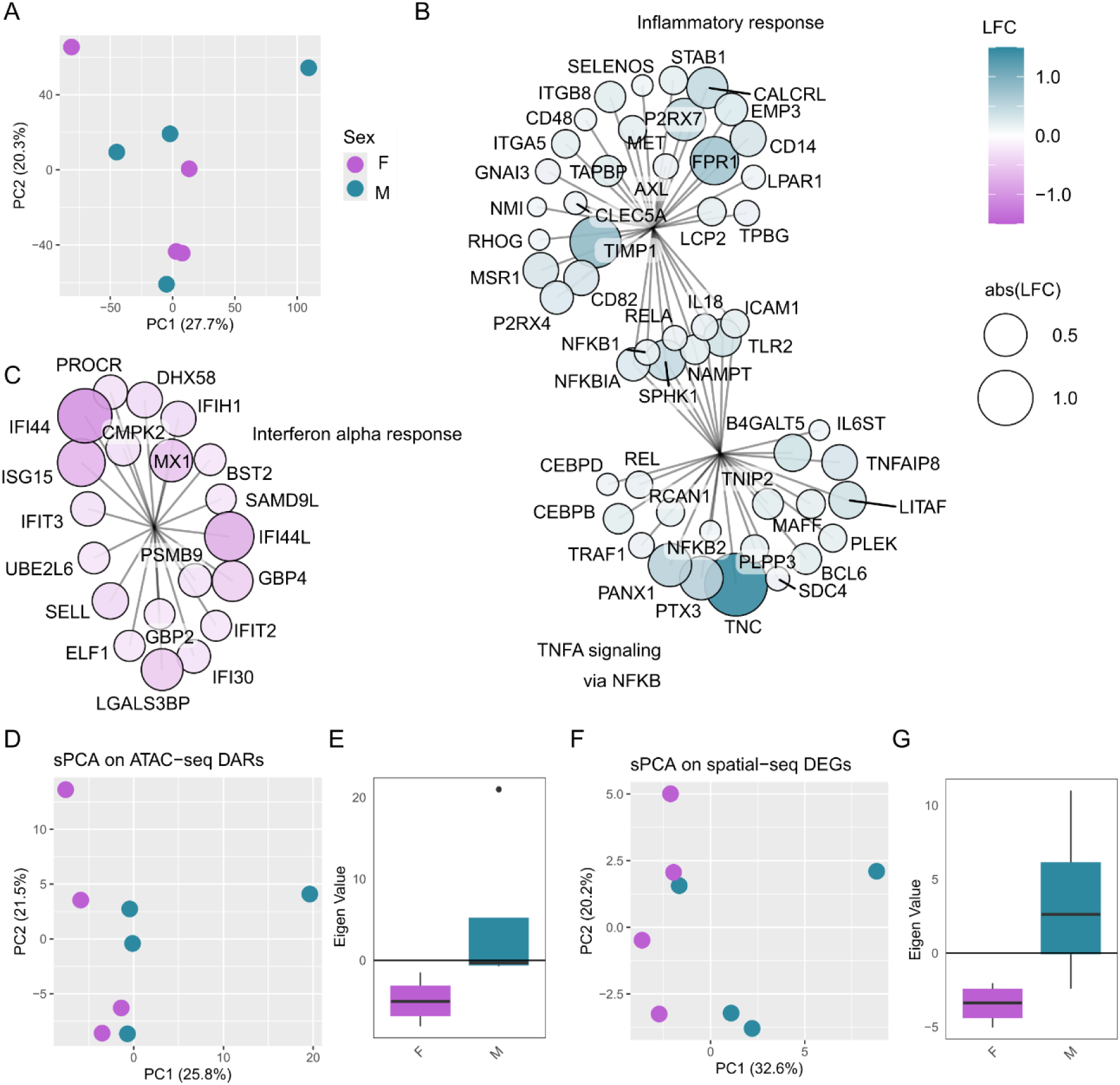
Sexual dimorphism in the hDRG. A. PCA of ganglia samples (merged by replicate). B-C. Representative Gene Set Enrichment Analysis (GSEA) pathway enrichments (male = positive, female = negative LFC). B. TNFα signalling via NFκB with overlap to Inflammatory response. C. IFNα response. Supervised PCA (sPCA) and corresponding eigen values (PC1) of ganglia samples on DARs (ATAC-seq, [25]) (D-E), and DEGs (RNA-seq, [55]). Related to SFig 4-7. n = 4M + 4F donor DRGs.

To see if there was a shared signature of sexual dimorphism across omics types (ie. ATAC-seq and RNA-seq), we next clustered the ganglia samples by supervised PCA (sPCA) against previously published differentially accessible regions (DARs) [25] and differentially expressed genes (DEGs, [55]) comparing hDRG of male and female donors (Fig 4D-G).

We see a separation of male and female proteomic samples by sPCA against sexually dimorphic DARs (ATAC-seq, Fig 4D-E) and DEGs (RNA-seq, Fig 4F-G), although this trend is not statistically significant (Welch’s, p ∼ 0.1) in either case. PCA and sPCA are dimensionality reduction techniques that calculate a covariance matrix to represent high-dimensional data (e.g., many proteins) in fewer dimensions (e.g., the first and second principal components), with the components optimized to maximize variance. From this, the contribution of each protein to each principal component can still be extracted. Here, eigengenes were extracted from the first principal component to see what was driving this separation by sex, with a number of proteins in the TNFα signalling pathway showing high loading values, suggesting they contribute to driving the difference across sex (SFig 5).

To see if the pathway changes from our proteomic data are shared across omics, we next performed GSEA against pseudobulk spatial ATAC-seq (SFig 6A), as well as bulk RNA-seq from thoracic vertebrectomy participants (SFig 6B) [25,45]. TNFα signalling via NFκB was enriched in males across datasets, suggesting a cross-omic pattern of sexual dimorphism.

The nerve root (SFig 7A) exhibits slightly fewer protein identifications (SFig 7B) than the ganglia, but again do not cluster by sex (SFig 7C) or show differentially expressed proteins by sex (SFig 7D, Supplemental Table 6-7). Mirroring the GSEA from Fig 4, we see a different pattern of sexual dimorphism in the nerve root compared to the ganglia. Like the ganglia, we see an enrichment for Oxidative Phosphorylation in female donors, but do not replicate the differences in IFNα Response or TNFα signalling via NFκB (SFig 7E). By sPCA, we also do not see a clear separation by sex using DARs (SFig 7F) or DEGs (SFig 7G) from the corresponding hDRG ATAC-seq and RNA-seq datasets [25,55].

### Sexual dimorphism across omics datasets

To probe shared patterns of sexual dimorphism across omics types, we used a multi-study factor analysis on available hDRG datasets (Fig 5, SFig 8) [25,45]. Multi-study factor analysis allows us to examine class separation (here, male and female) through shared factors across omics types, where each relevant factor has interpretable underlying constructs.

**Figure 5.**
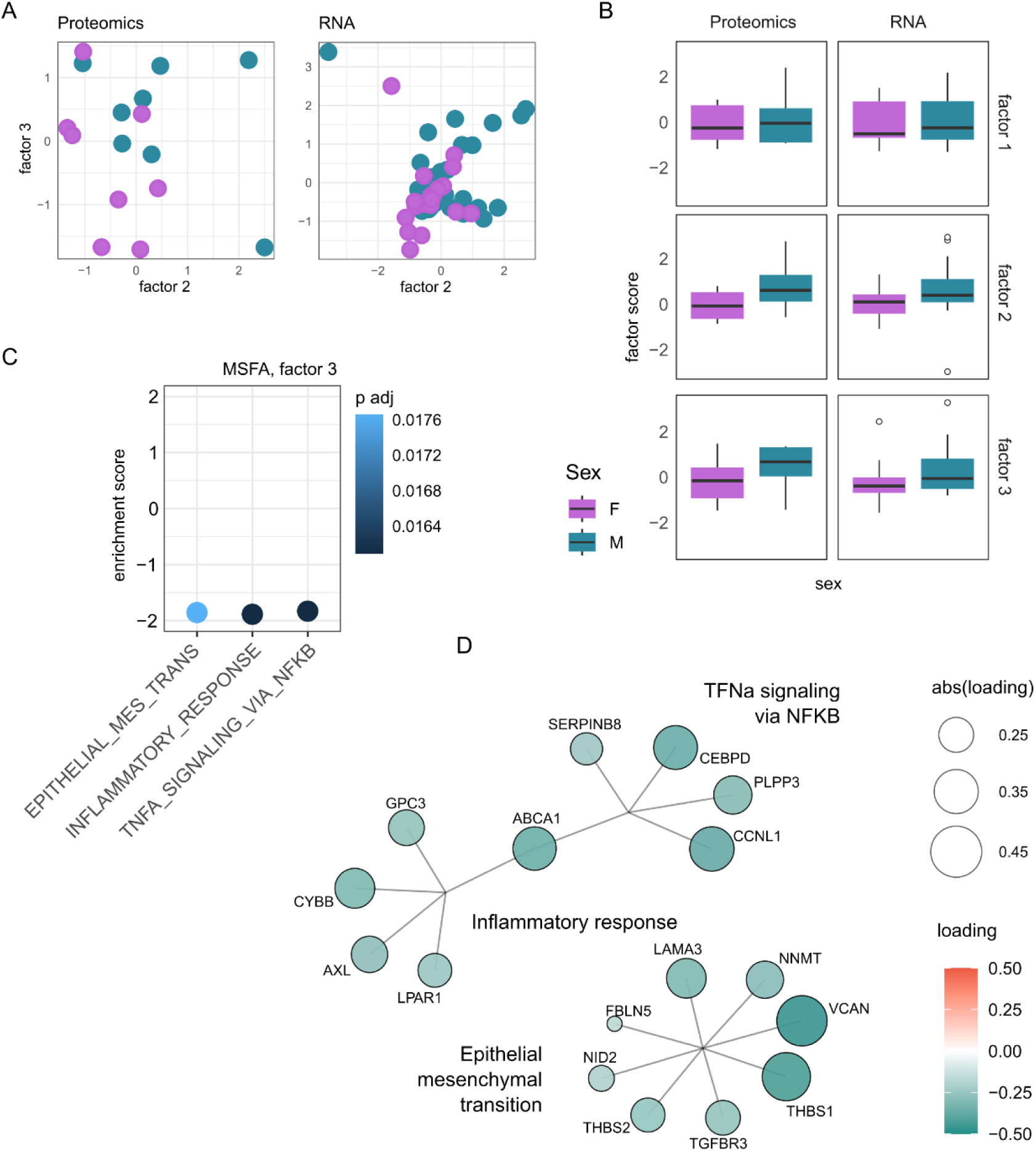
Multi-study factor analysis between bulk RNA-seq and bulk proteomic data, based on differentially accessible regions from bulk ATAC-seq. A. Scatter plot by factor, for proteomics (left) and transcriptomics (right). B. Boxplot of Factor scores, stratified by sex, bars = ±1.5*interquartile range. C. GSEA on factor 3 (ranked loadings). D. Pathways + gene loadings from Gene Set Enrichment Analysis (GSEA) output of Factor 3. Related to SFig 8.

We used the most comprehensive bulk RNA-seq dataset available to date, looking at neuropathic pain in a participant cohort of thoracic vertebrectomy patients undergoing surgery. Of the 70 DRG sequenced, 50 showed neuronal enrichment, as described in the original publication [45]. These 50 DRG, removed during thoracic vertebrectomy surgical procedure, were stratified by sex for comparison to our hDRG proteomic dataset (both ganglia and nerve root). There were too few samples from the hDRG ATAC-seq datasets for a factor analysis, thus we instead restricted our factor analysis to gene/proteins which correspond to DARs between male and female donors (see methodology) [25].

Likely because of the small sample sizes, only 3 shared factors were extracted, with Factors 2-3 both showing a separation by sex (Fig 5A-B). GSEA against the ranked loading revealed no hallmark pathway enrichment for Factor 2, but 3 enriched pathways for Factor 3: TNFα signalling via NFκB, Inflammatory Response, and Epithelial Mesenchymal Transition (Fig 5C-D, SFig 8A). While there are overlapping terms driving TNFα signalling via NFκB and Inflammatory Response, the enrichment of Epithelial Mesenchymal Transition is independent of this (Fig 5D), but was also detected in our proteomics data (SFig 4B), and in bulk RNA-seq (SFig 6B).

### TNFα release from hDRG immune cells

TNFα signalling consistently appears as a sexually dimorphic pathway in these omics datasets, spanning multiple sets of donors and a large cohort of surgical participants.

If the sexual dimorphism we see is functional relevant, we would expect a few things: Firstly, drugs targeting the pathway should show response differences in men and women (they do, Table 2). Secondly, SNPs against core genes in the pathway would be related to pain and sensory processing (they are, Table 3). Here, sexual dimorphism at the genetic level is also discussed for the TNFα promotor in the context of migraine Fawzi et al., (2015).

**Table 2.** Example clinical outcomes on TNFα inhibitors and sexual dimorphism.

| PMID | Citation | Title | Condition | Relevant outcome |
| --- | --- | --- | --- | --- |
| *Primary examples* |  |  |  |  |
| PMID: 19815494 | [10] | Effectiveness of adalimumab in treating patients with active psoriatic arthritis and predictors of good clinical responses for arthritis, skin and nail lesions | Psoriatic arthritis | Male sex increased the chance of achieving a good clinical response |
| PMID: 28762060 | [12] | A 2-year observational study on treatment targets in psoriatic arthritis patients treated with TNF inhibitors | Psoriatic arthritis | Females less likely to achieve remission |
| PMID: 16705046 | [31] | Predictors of response to anti-TNF-α therapy among patients with rheumatoid arthritis: results from the British Society for Rheumatology Biologics Register | RA | Females less likely to achieve remission |
| PMID: 23322995 | [63] | Sex-dimorphic adverse drug reactions to immune suppressive agents in inflammatory bowel disease | IBD | Higher ADR in women taking TNFα-inhibitor |
| PMID: 31310690 | [49] | Earlier discontinuation of TNF-α inhibitor therapy in female patients with inflammatory bowel disease is related to a greater risk of side effects | IBD | Higher ADR in women taking TNFα-inhibitor |
| PMID: 12492735 | [14] | Infusion reactions to infliximab in children and adolescents: frequency, outcome and a predictive model | Crohn's | Female sex as a predictor for side effects in TNFα-inhibitors for Crohn's |
| *Example Reviews* |  |  |  |  |
| PMID: 37820857 | [13] | Sex-oriented perspectives in immunopharmacology | Mixed | TNFis show "higher efficacy and adherence in males, More frequent serious infections in males, more frequent toxic liver disease and lupus-like syndrome in women" |
| PMID: 29754330 | [47] | Gender Differences in Axial Spondyloarthritis: Women Are Not So Lucky | Axial Spondylo-arthritis | Increased TNF levels in only male AS patients, treatment efficacy of TNFi is significantly lower in women compared to men with axSpA, and they have a significantly lower drug adherence |
| PMID: 26490106 | [53] | Rate of discontinuation and drug survival of biologic therapies in rheumatoid arthritis: a systematic review and meta-analysis of drug registries and health care databases | RA | Female sex as a predictor of discontinuation |

RA: Rheumatoid Arthritis, IBD: Inflammatory Bowel Disease

**Table 3.**
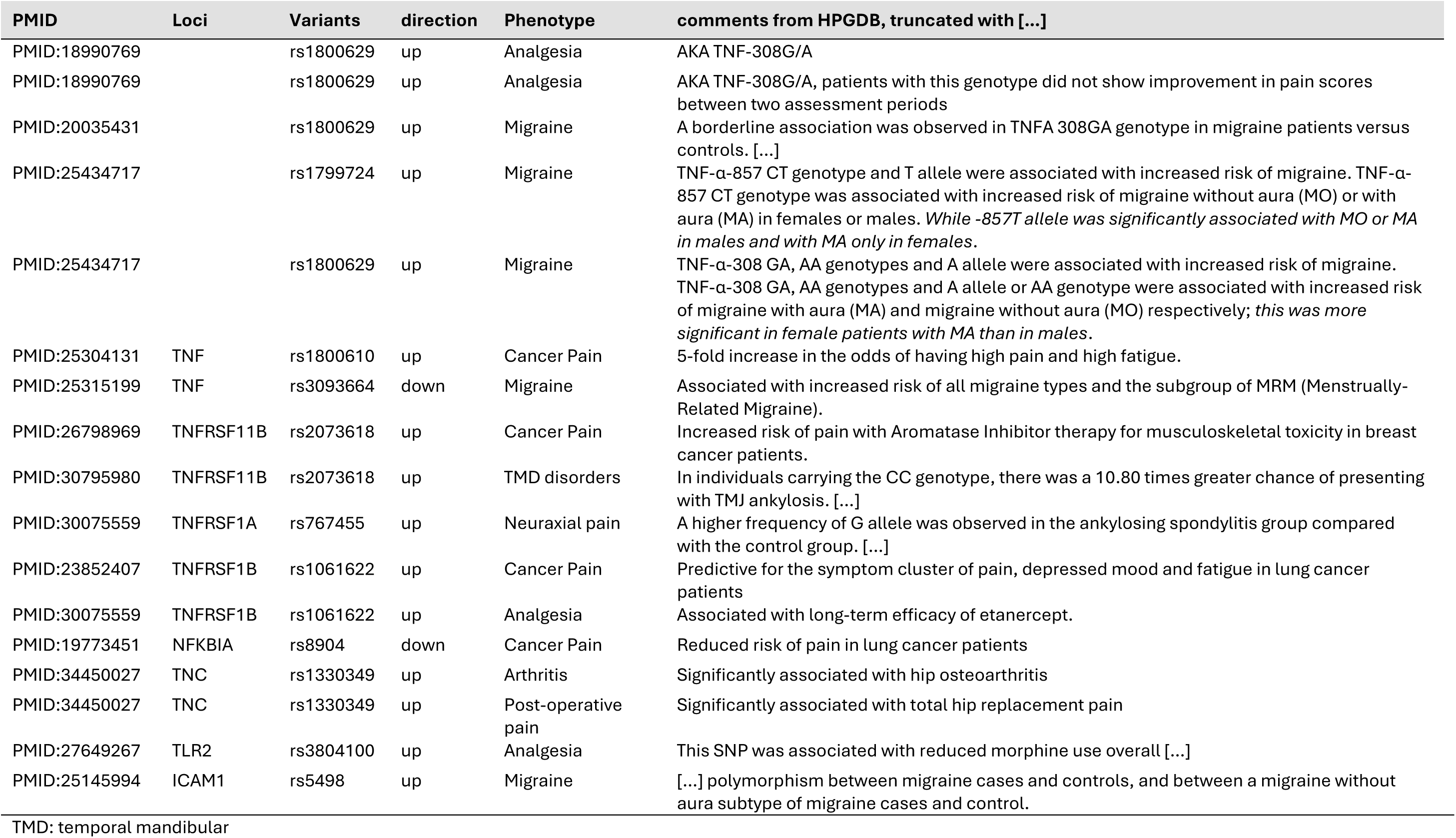
Summary of pain-related GWAS hits for the TNFα signalling pathway from the HPGDB [40].

| PMID | Loci | Variants | direction | Phenotype | comments from HPGDB, truncated with [...] |
| --- | --- | --- | --- | --- | --- |
| PMID:18990769 |  | rs1800629 | up | Analgesia | AKA TNF-308G/A |
| PMID:18990769 |  | rs1800629 | up | Analgesia | AKA TNF-308G/A, patients with this genotype did not show improvement in pain scores between two assessment periods |
| PMID:20035431 |  | rs1800629 | up | Migraine | A borderline association was observed in TNFA 308GA genotype in migraine patients versus controls. [...] |
| PMID:25434717 | | rs1799724 | up | Migraine | TNF-α-857 CT genotype and T allele were associated with increased risk of migraine. TNF-α-857 CT genotype was associated with increased risk of migraine without aura (MO) or with aura (MA) in females or males. *While -857T allele was significantly associated with MO or MA in males and with MA only in females.* |
| PMID:25434717 | | rs1800629 | up | Migraine | TNF-α-308 GA, AA genotypes and A allele were associated with increased risk of migraine. TNF-α-308 GA, AA genotypes and A allele or AA genotype were associated with increased risk of migraine with aura (MA) and migraine without aura (MO) respectively; *this was more significant in female patients with MA than in males.* |
| PMID:25304131 | TNF | rs1800610 | up | Cancer Pain | 5-fold increase in the odds of having high pain and high fatigue. |
| PMID:25315199 | TNF | rs3093664 | down | Migraine | Associated with increased risk of all migraine types and the subgroup of MRM (Menstrually-Related Migraine). |
| PMID:26798969 | TNFRSF11B | rs2073618 | up | Cancer Pain | Increased risk of pain with Aromatase Inhibitor therapy for musculoskeletal toxicity in breast cancer patients. |
| PMID:30795980 | TNFRSF11B | rs2073618 | up | TMD disorders | In individuals carrying the CC genotype, there was a 10.80 times greater chance of presenting with TMJ ankylosis. [...] |
| PMID:30075559 | TNFRSF1A | rs767455 | up | Neuraxial pain | A higher frequency of G allele was observed in the ankylosing spondylitis group compared with the control group. [...] |
| PMID:23852407 | TNFRSF1B | rs1061622 | up | Cancer Pain | Predictive for the symptom cluster of pain, depressed mood and fatigue in lung cancer patients |
| PMID:30075559 | TNFRSF1B | rs1061622 | up | Analgesia | Associated with long-term efficacy of etanercept. |
| PMID:19773451 | NFKBIA | rs8904 | down | Cancer Pain | Reduced risk of pain in lung cancer patients |
| PMID:34450027 | TNC | rs1330349 | up | Arthritis | Significantly associated with hip osteoarthritis |
| PMID:34450027 | TNC | rs1330349 | up | Post-operative pain | Significantly associated with total hip replacement pain |
| PMID:27649267 | TLR2 | rs3804100 | up | Analgesia | This SNP was associated with reduced morphine use overall [...] |
| PMID:25145994 | ICAM1 | rs5498 | up | Migraine | [...] polymorphism between migraine cases and controls, and between a migraine without aura subtype of migraine cases and control. |

TMD: temporal mandibular

To further support the relevance of TNFα sexual dimorphism in the peripheral nervous system, we investigated TNFα release and signalling in the hDRG directly (Fig 6).

**Figure 6.**
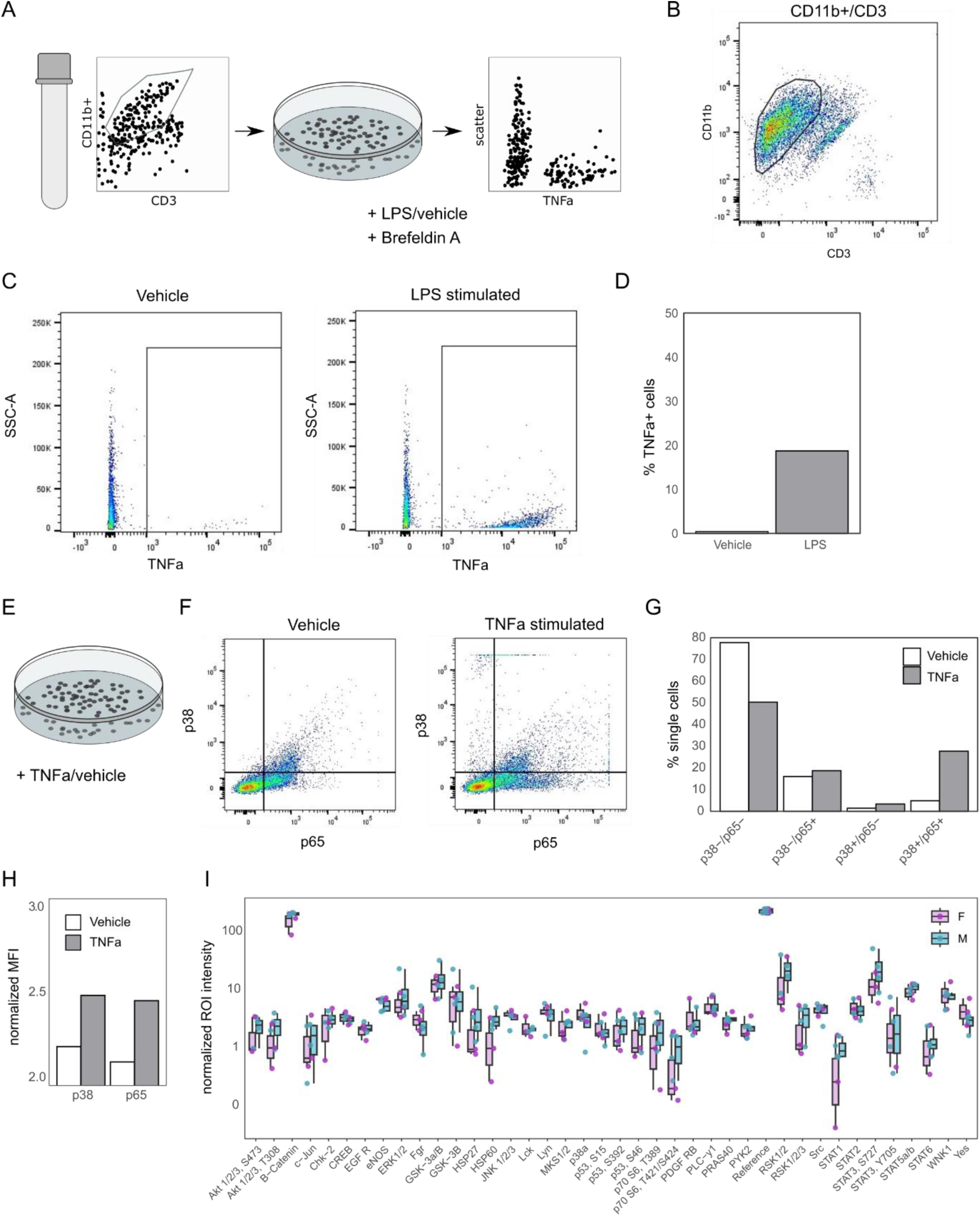
TNFα signalling in the hDRG. A. FACS experiment overview, for isolation and culturing of myeloid cells followed by stimulation *in vitro* (n = 1 donor, Male). B. Myeloid cell gating strategy, full strategy shown in SFigure 9. C. Representative flow cytometry for Vehicle (left) and LPS (right) stimulation of myeloid cells. D. Percentage of TNFα+ single cells after LPS stimulation. E. Schematic for TNFα stimulation of myeloid cells. F. Transcription factor phosphorylation (p38 and p65) from Vehicle (left) and TNFα (right) treated cells. G. Quantification of (F), % of single cells per condition/gate. H. Normalized Mean Fluorescent Intensity (MFI) for p38 and p65 per condition. I. Phosphorylation array of Male (M) and Female (F) hDRG protein samples (anova: Sex (p = 0.0266), Sex:Site interaction (p = 2e-06), n = 4 per group). Related to SFigure 9-11.

To determine if myeloid immune cells from DRG parenchyma could be a local source of TNFα, we isolated CD11b+ cells from hDRG via FACS (Fig 6A-B). Following isolation, cells were stimulated with LPS (10 ng/mL) or vehicle (RPMI media) with or without the intracellular protein transport inhibitor Brefeldin A for 16 hours. In myeloid cells without Brefeldin A, we found a robust increase in levels of TNFα following LPS stimulation (SFigure 10, n=5). The statistical power to test sexual dimorphism (assuming a 2-fold effect size, alpha = 0.1 and n=2-3) is 0.12-0.14 with a two-sided t-test (’ pwr’ package, in R). Given this limited power, we are unable to robustly assess male/female differences based on the variability in the current data.

We complemented this data with myeloid cells co-stimulated with Brefeldin A followed by fixation, permeabilization, and staining for intracellular TNFα production with a TNFα antibody (Fig 6C-D). Following LPS stimulation, there was a higher percentage of cells with TNFα staining (18.7%) when compared to vehicle treated cells (0.36%) (Fig 6C-D).Together, these findings suggest that hDRG myeloid cells can serve as a local source of TNFα production.

### Phosphorylation in the hDRG

In parallel, we examined if TNFα stimulation could drive increases in the phosphorylation of transcription factors p38 and p65. CD11b+ cells were stimulated with TNFα (10ng/mL) or its vehicle (1X PBS) for 30 minutes (Fig 6E). Cells were then collected, fixed, permeabilized, and stained with p38 and p65 antibodies. We observed a higher percentage of CD11b+ cells stained for p38 and p65 upon TNFα stimulation when compared with vehicle treated cells (Fig 6F-G). TNFα treated cells also exhibited a higher median fluorescent intensity signal of both p38 and p65 antibodies (Fig 6H), together suggesting that TNFα can drive the phosphorylation of p38 and p65 in the DRG.

Finally, we asked whether phosphorylation in general may be sexually dimorphic (indirectly of TNFα stimulation) in hDRG. We assessed the phosphorylation status of kinases involved in prominent cellular signalling pathways (e.g. STAT, EGF, ERK1/2) in matched protein samples from our proteomic experiment (4M + 4F) to query sex-specific phosphorylation differences in our pain-free donors (Fig 6I). We see significant differences in Sex (p = 0.0266), and the Sex:Site interaction (p = 2e-06), but we do not see differences for individual phosphorylation sites (Tukey’s honestly significant difference (Tukey’s HSD), adjusted p ∼ 1 for all sites).

## Discussion

This study explores sexual dimorphism in a proteomics dataset of the hDRG. It complements previous ATAC-seq and RNA-seq work, and with deep coverage spanning over 12500 Gene Groups (and ∼ 9000 unique proteins) per tissue it provides the most comprehensive proteomic data to date for the hDRG. Despite our small sample size, we report distinct sex differences with a particular focus on TNFα signalling, highlighting that these differences are seen consistently across cohorts and omics technologies. We have paired this to *in vitro* work on freshly isolated hDRG immune cells, confirming the functional relevance of this pathway in the hDRG. Collectively, this indicates a broad sexual dimorphism in the hDRG, with likely functional and clinical implications.

In addition to providing the first human-centric DIA-MS proteomics reference of the hDRG, we also improve the coverage over pre-clinical DRG proteomes in terms of ion channel and membrane protein detection [6]. In the current study, we highlight membrane proteins relevant to pain- and neuroimmune- conditions, even detecting large membrane proteins like PIEZO1.

We validate this dataset across the nerve root and ganglia, highlighting differences in ion channel expression and myelin-related proteins by tissue. In the nerve root, we see an enrichment of terms related to myelination via ‘myelination’, ‘axon ensheathment’, and ‘gliogenesis’, among others, while also observing an enrichment in ion regulation, including ‘potassium ion homeostasis’, ‘intracellular sodium ion homeostasis’, and ‘sodium ion transmembrane transport’. Fewer GO terms are enriched in the ganglia, these are related to ‘tRNA aminoacylation’ and ‘amino acid activation’ highlighting the role of cell bodies in translation.

While both the nerve and ganglia contain primary afferent components (soma vs axons), mRNAs can be transported from the soma for localized translation in the axons [48]. These tissues also contain different proportions of non-neuronal cell types, with satellite glial cells wrapping around soma and Schwann cells (both myelinating and non-myelinating) playing essential roles in axonal integrity. As a result, peripheral nerves and ganglia are uniquely affected in conditions such as diabetic and chemotherapy-induced peripheral neuropathies, and knowledge of their protein-setup can lay a foundation for future research.

When probing sexual dimorphism in the ganglia and nerve root, we see a shared pattern of ‘Oxidative Phosphorylation’ enrichment in female donors, as well as differences in ‘TNFα’ and ‘IFNα’ signalling. Notably, we see TNFα signalling enriched in male ganglia across datasets, spanning from different accessible gene regions (via ATAC-seq, [25]) to different transcriptional profiles in human participants [45]. By detecting this sexual dimorphism across omics types, as well as across samples from organ donors and human participants, we gain confidence in this being a true biological signal. Paired to this, there is strong evidence from clinical trials that TNFα inhibitors affect men and women differently [13], suggesting a meaningful molecular signature.

In mice, TNFα signalling in the DRG was recently shown to be sexually dimorphic in an experimental autoimmune encephalomyelitis model [39]. In humans, this pathway is frequently studied in circulating immune cells, and TNFα signalling here can be regulated by testosterone levels [37]. Using FACS/flow cytometry on freshly isolated hDRG, we first examine the functional relevance of this pathway in the hDRG specifically: When stimulated, CD11b+ cells from hDRG release TNFα, and TNFα stimulation can alter phosphorylation levels.

Using a targeted phosphorylation approach, we then investigate sexual dimorphism at the level of post-translational modifications. Here, we see an interaction of sex and phosphorylation site in the hDRG. Together this provides an initial avenue for follow-up in humans, as previous work in mice by Maguire and colleagues report phosphorylation differences in the context of the TNFα signalling pathway dimorphism [39].

## Limitations

This dataset presents the highest proteome coverage of the hDRG to date and overcomes many of the known difficulties in detecting membrane proteins, but is not complete. We observe ∼20% of ion channels with detectable transcripts in bulk RNA-seq data, and we detect less than 10% of the corresponding GPCRs detected in a comparable bulk RNA-seq dataset [45]. Prior studies highlight functional expression of membrane proteins like ASICs, opioid receptors (OPRs), and PIEZO2 in DRG which are missing here, thus the lack of detection is likely technical (given their low abundance), opposed to biological. Increases in sample sizes, paired to further improvements in sensitivity and sample preparation protocols will facilitatethe detection of very low abundant membrane proteins in complex protein mixtures in future work [44,60].

Proximal and distal nerve regions exhibit prominent differences and are uniquely affected by length-dependent pathologies (eg. diabetic peripheral neuropathy [22]), thus our nerve data likely do not represent the proteomes of distal nerves or terminal endings. As technology develops, added information on peripheral endings in the skin and viscera will also be beneficial, as current proteomics of human skin does not include common neuronal markers [21].

## Future Directions

The full implications of the sexual dimorphism reported here remain unclear: previous work in healthy trans men show that testosterone (as hormone replacement therapy, HRT) can shift immune function in plasma samples in line with the dimorphism of the hDRG we report here [37]. Within 12 months of starting HRT, these men show increased TNF signalling through NFκB, paired to a decrease in the IFNα response, highlighting adaptations to immune function in response to increasing testosterone levels.

Beyond TNFα signalling, varying levels of testosterone in cis men correlate to immune response after vaccination [26], and HRT in post-menopausal women can elicit immune changes as early as 1 month [35]. Hormone differences are highly relevant in autoimmune conditions as well: while HRT in menopausal women may increase the risk of late-onset RA, hormonal oral contraceptives have been shown to reduce the risk of RA in women [29].

Given the variation across humans, and the adaptive nature of the immune system, the differences reported likely exist on a spectrum which can change with medication and age. In the hDRG specifically, the functional implications require more research. We show that TNFα is released from myeloid cells from DRG parenchyma, and that TNFα stimulation of hDRG immune cells can result in protein phosphorylation, but a direct causation to sexually dimorphic PTMs - and any downstream functional changes - is still needed.

Moving forward, differing immune signatures in their local environment should be accounted for in future studies, especially when considering developmental markers like puberty and menopause.

This comprehensive, human-centric proteomic dataset complements previously published ATAC-seq and RNA-seq data, providing a quantitative reference proteome of the hDRG. Data are searchable at https://sensoryomics.com/.

Using tissue from both human participants and donors, we show evidence for sexual dimorphism in TNFα signalling spanning the epigenetic to proteomic signature. Taken together, these results underscore a relevant sexual dimorphism in the hDRG, with significant implications for sensory and pain-related translation.

## Code Availability

Analysis scripts are available at https://github.com/aliibarry/hDRG-proteomics.

## Supporting information

STable 9

STable 10

STable 11

STable 1

STable 2

STable 3

STable 4

STable 5

STable 6

STable 7

STable 8

## Data Availability

Proteomic data (raw and processed) have been deposited via SPARC, DOI:10.26275/z7uy-kuif [4], and have been added to a searchable database at https://sensoryomics.com. Raw data have also been deposited to the ProteomeXchange Consortium via the PRIDE partner repository [43] with the dataset identifier PXD061129. Corresponding metadata and processed expression matrix are also on github, under ‘data/processed/*’, and for ease of access the proteomic expression matrix is also included here as Supplemental Table 11. Source data for the targeted phosphorylation array are in Supplemental Table 9; imaged membranes and selected ROIs are shown in Supplemental Figure 11.

## Acknowledgements

The authors thank the organ donors and their families for their gift of life. Further thanks to E. Vines and the staff at the Southwest Transplant Alliance for coordinating DRG recovery from organ donation surgeries, P. Horton, G. Funk, and A. Cervantes at the Southwest Transplant Alliance for the organ donation surgeries where the DRG recovery was done, and the members of the Price laboratory, for assistance in DRG recovery. We thank members of the Schmidt laboratory for fruitful discussions.

This work was funded in part by National Institutes of Health grant U19NS130608 (T.J.P.), by the Austrian Science Fund (FWF) [10.55776/P36554] (M.S) and by the University of Vienna (M.S). The computational results presented have been achieved in part using the Vienna Scientific Cluster (VSC).

## Supplemental Tables

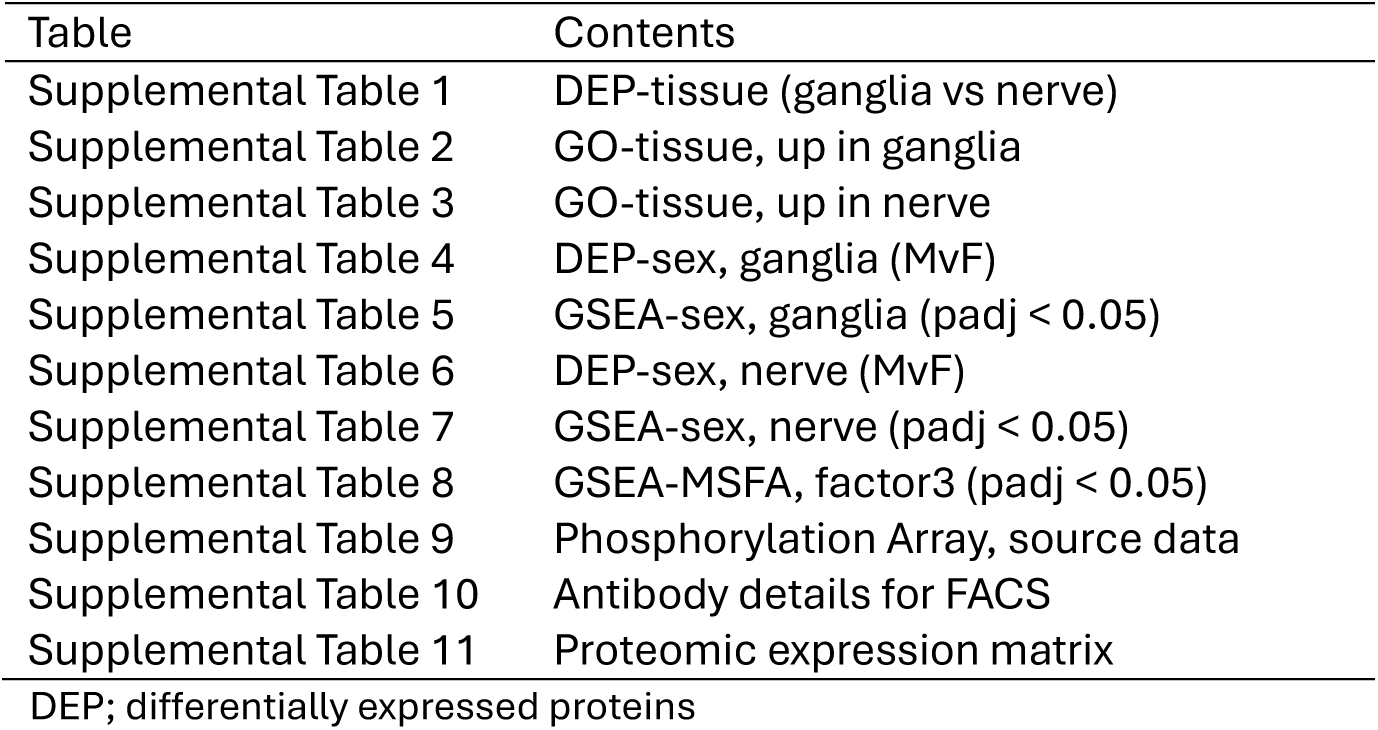

## Supplemental Figures

**SFigure 1.**
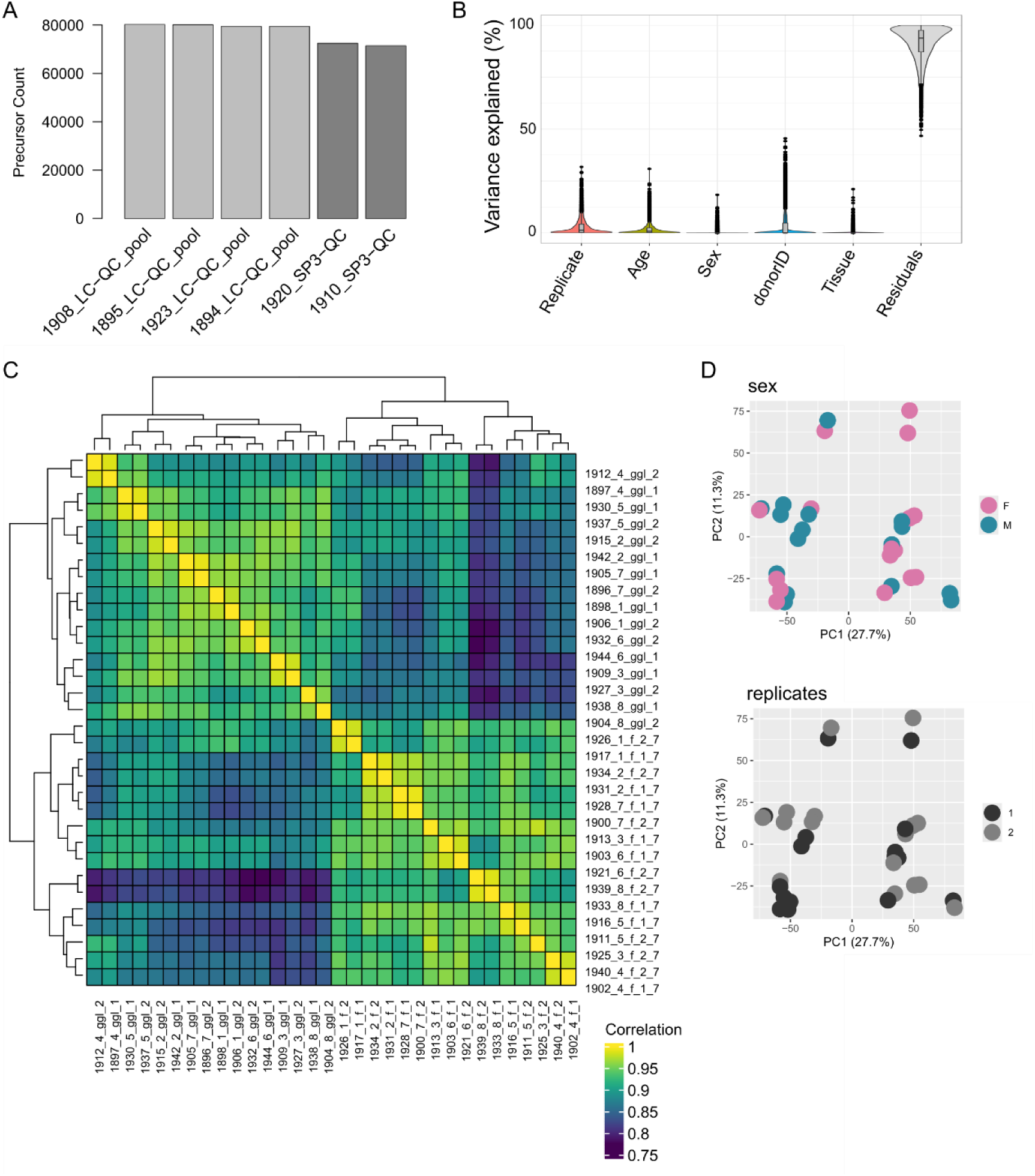
Quality control of the hDRG DIA-PASEF dataset. A. Precursors (q < 0.01) across reference quality control samples. B. Variance partitioning across all biological samples. C. Correlation across all biological samples (replicates as _1 or _2). D. PCA by sex and replicate for unmerged technical replicate samples.

**SFigure 2.**
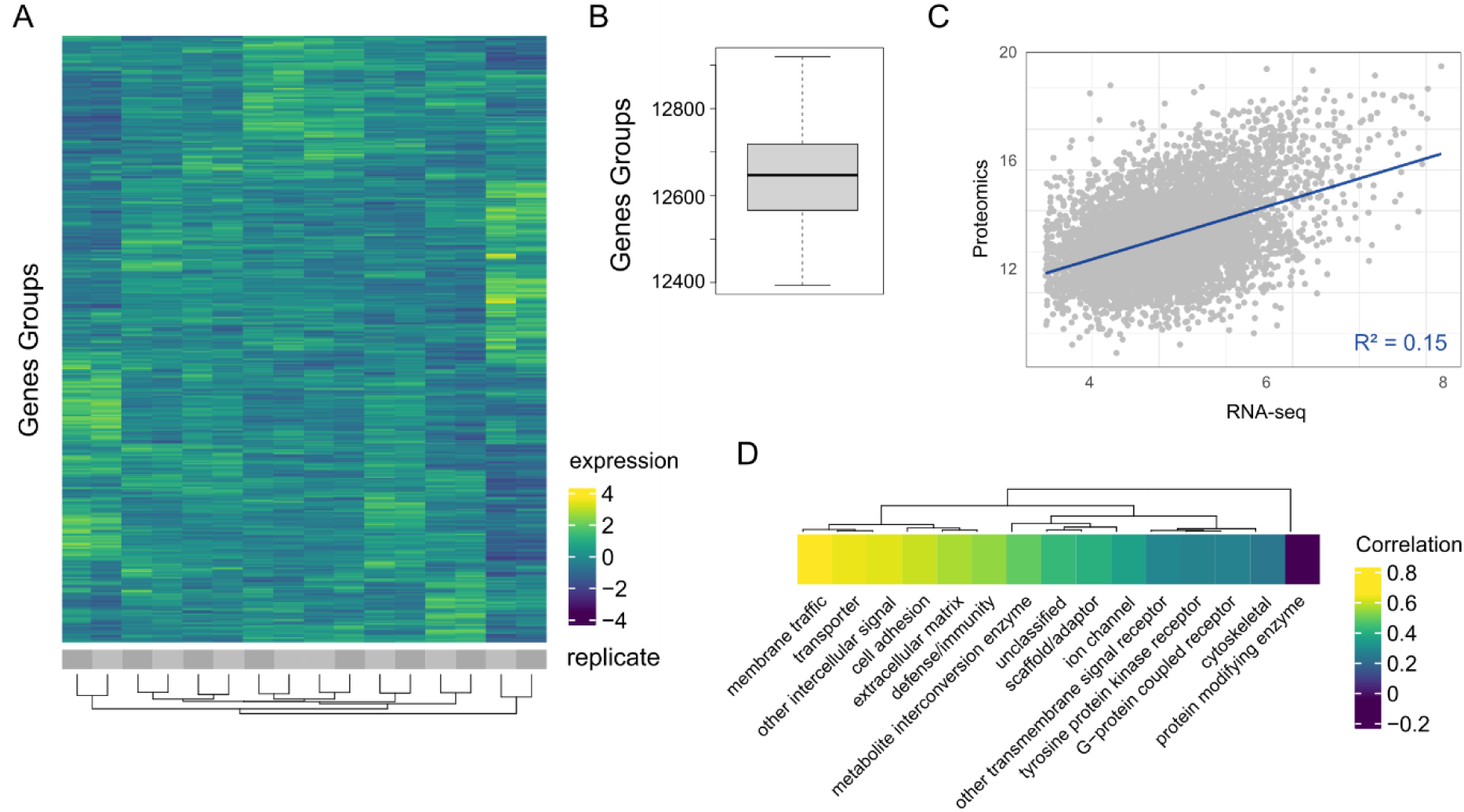
Quality control for the ganglia. A. Gene Group expression across replicates. B. Gene Group counts per sample. C. Correlation with hDRG RNA-seq data (average protein expression and average quantile-normalized transcripts per million (qnTPM)). D. Correlation with hDRG RNA-seq data separated by receptor group (min 15 proteins per group).

**SFigure 3.**
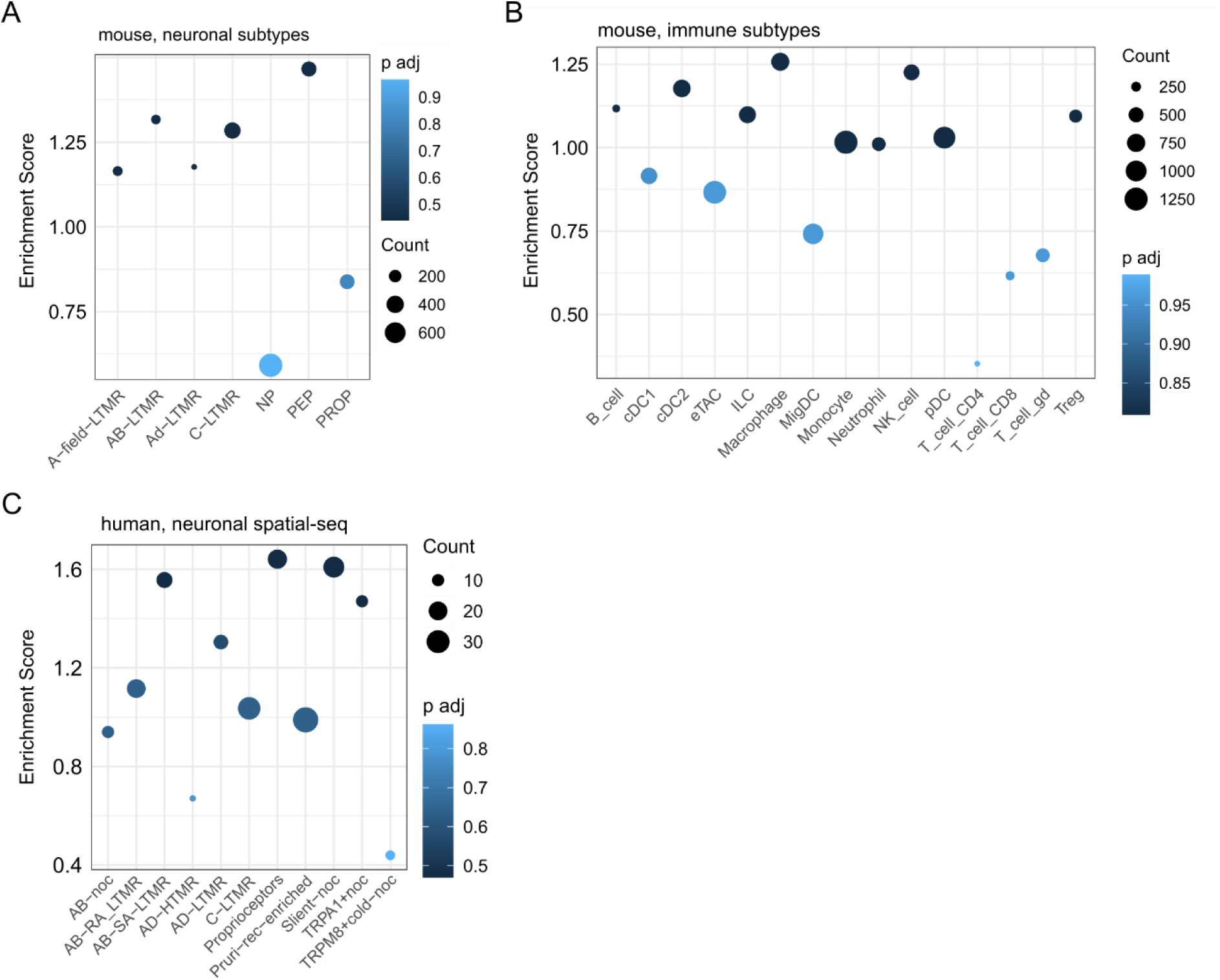
Gene set enrichment analysis (GSEA) for: A. mouse, neuronal. B. mouse, immune cells. C. human, neuronal gene sets against the hDRG proteome.

**SFigure 4.**
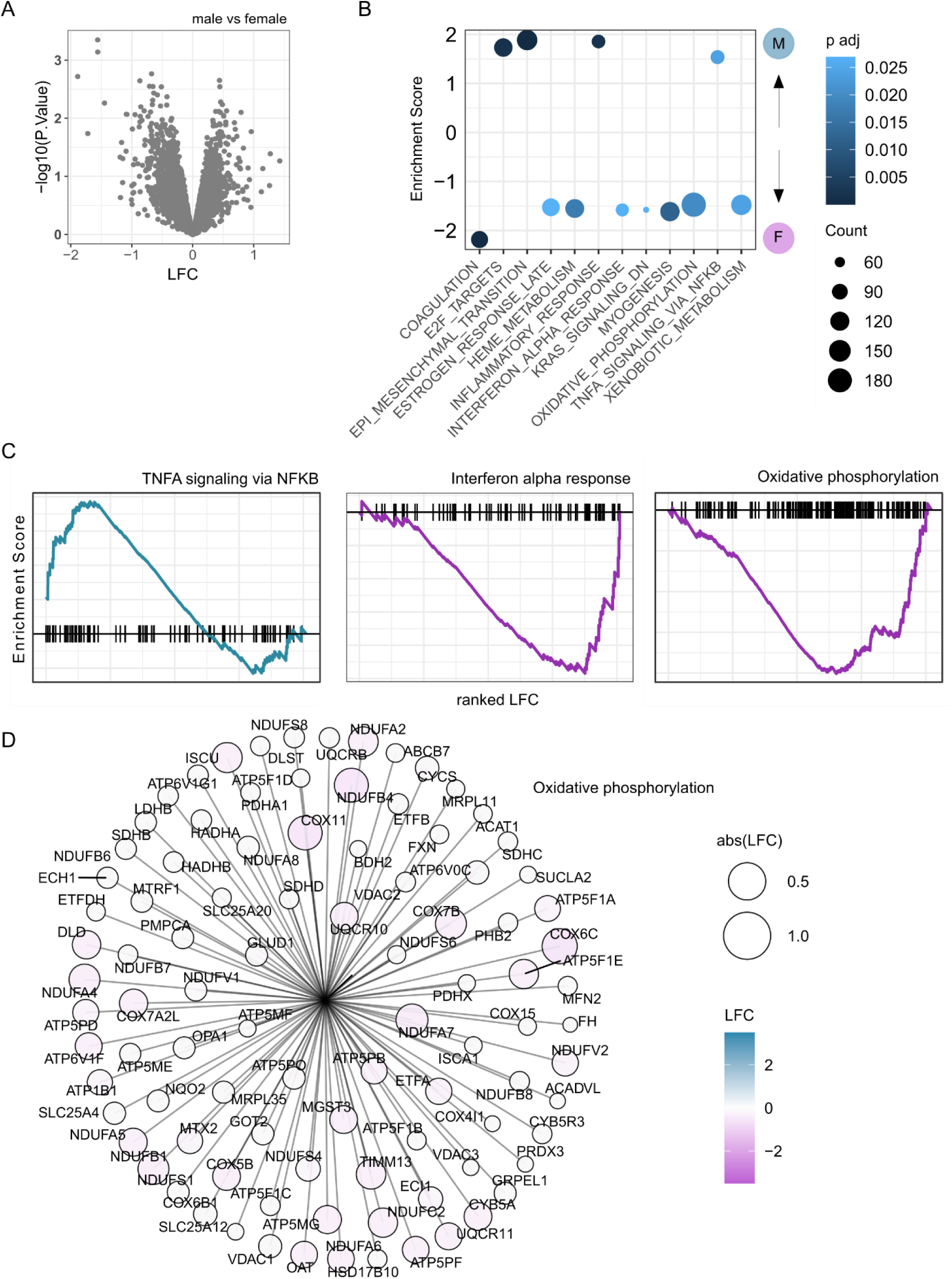
Sexual dimorphism in the hDRG. A. Volcano plot of DEPs. B. GSEA against hallmark pathways, males = positive. C. Representative gene rankings from B for pathways of interest. D. LFC for proteins in the ‘Oxidative Phosphorylation’ pathway. LFC = log fold change.

**SFigure 5.**
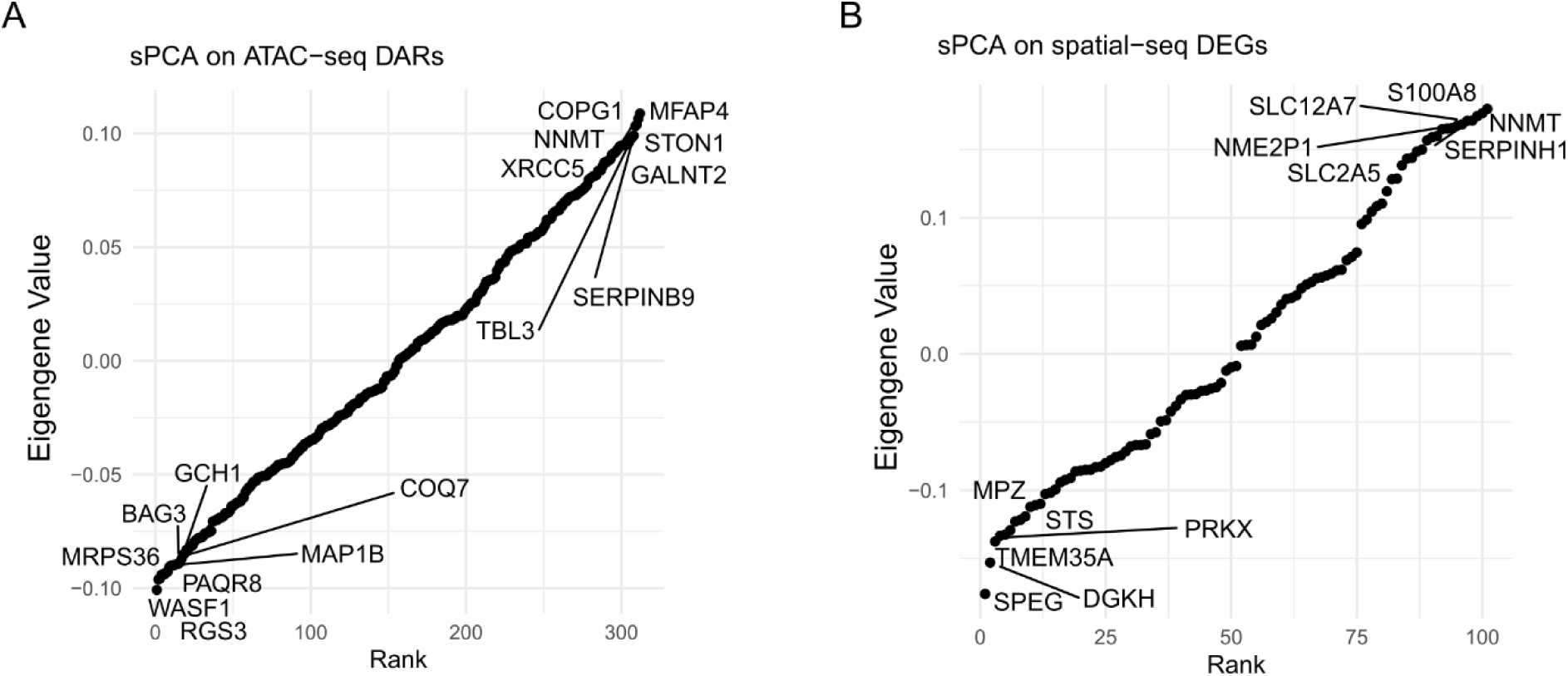
sPCA eigengenes from A. ATAC-seq DARs and B. spatial-seq DEGs, with tails labelled.

**SFigure 6.**
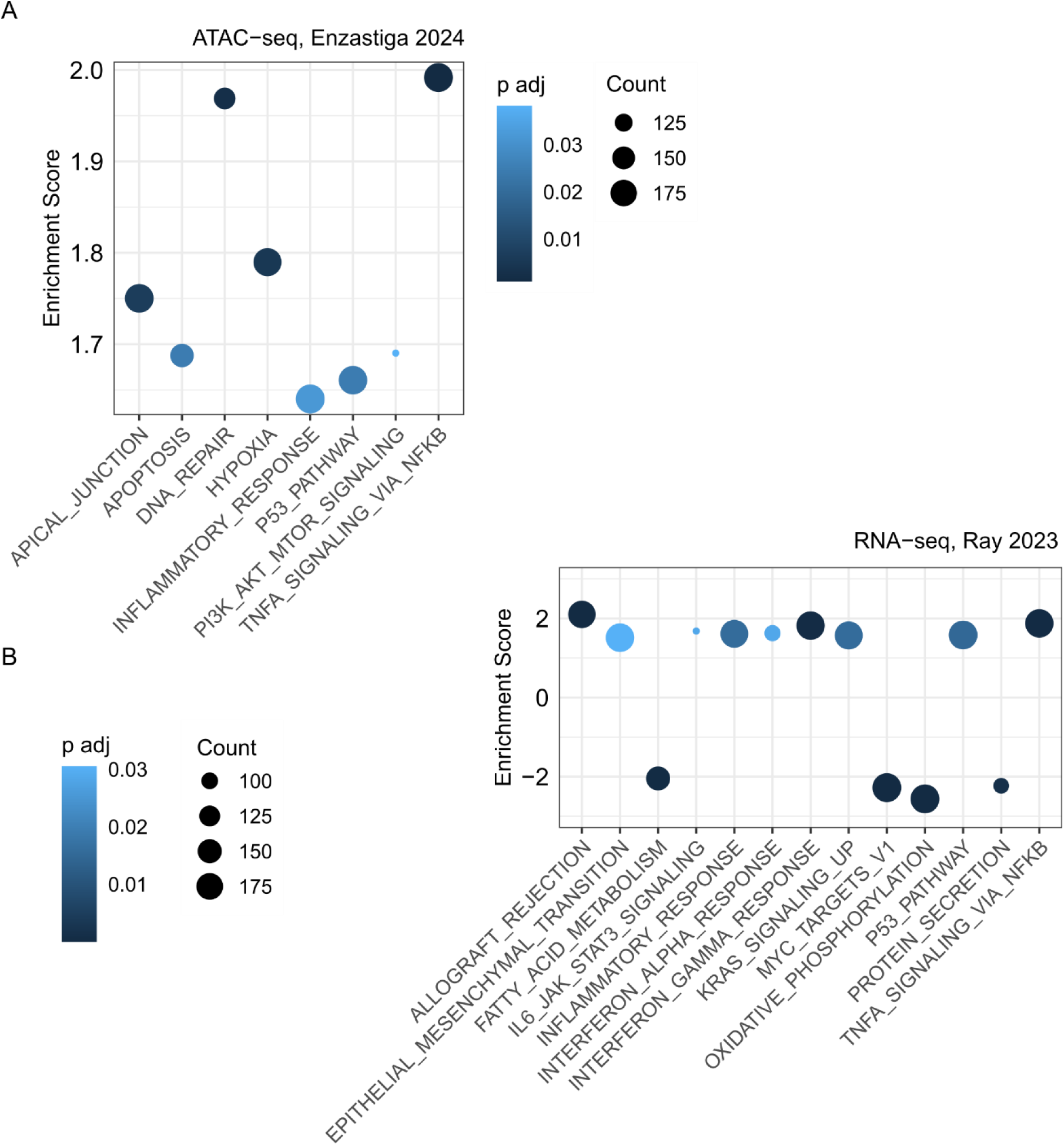
GSEA against previously published ATAC-seq and RNA-seq datasets [21,37]. A. ATAC-seq on donor samples, LFC for pseudobulk ATAC-seq spatial data. B. RNA-seq on participant samples.

**SFigure 7.**
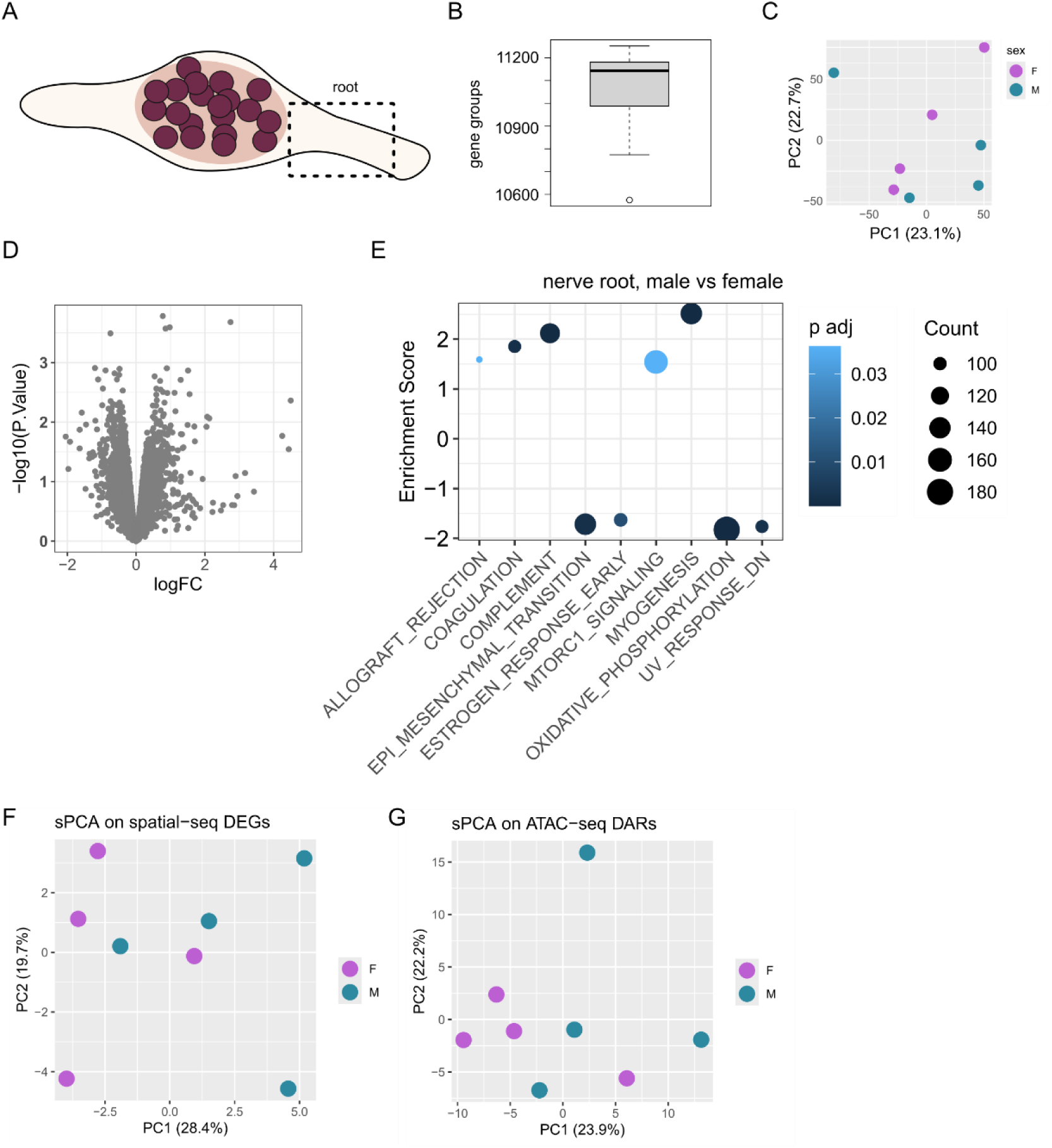
Proteome of the nerve root. A. Schematic. B. Gene Groups per sample. C. PCA by sex. D. Volcano plot, differential expression testing with limma, male = positive. E. GSEA against Hallmark pathways, male enrichment = positive. F-G. supervised PCA (sPCA) on differentially expressed genes (DEGs, F) and regions (DARs, G) from previously published reports on hDRG sexual dimorphism.

**SFigure 8.**
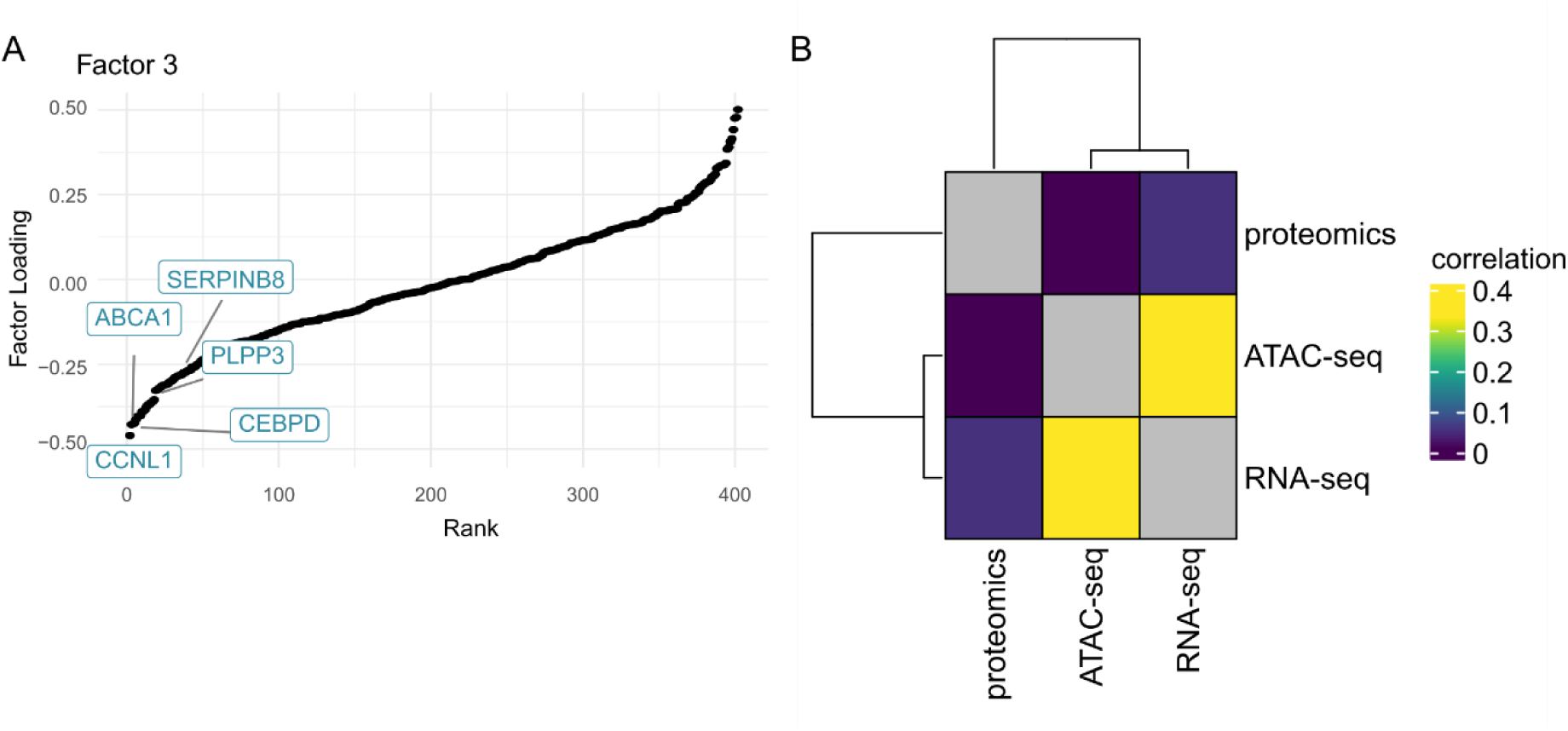
Multi-study factor analysis (MSFA) across omics datasets. A. Ranked factor loadings from Factor 3, with terms associated with TNFα signalling highlighted in the tail. B. Correlation between LFC for matched accessible regions, genes, and proteins, LFC for pseudobulk ATAC-seq spatial data.

**SFigure 9.**
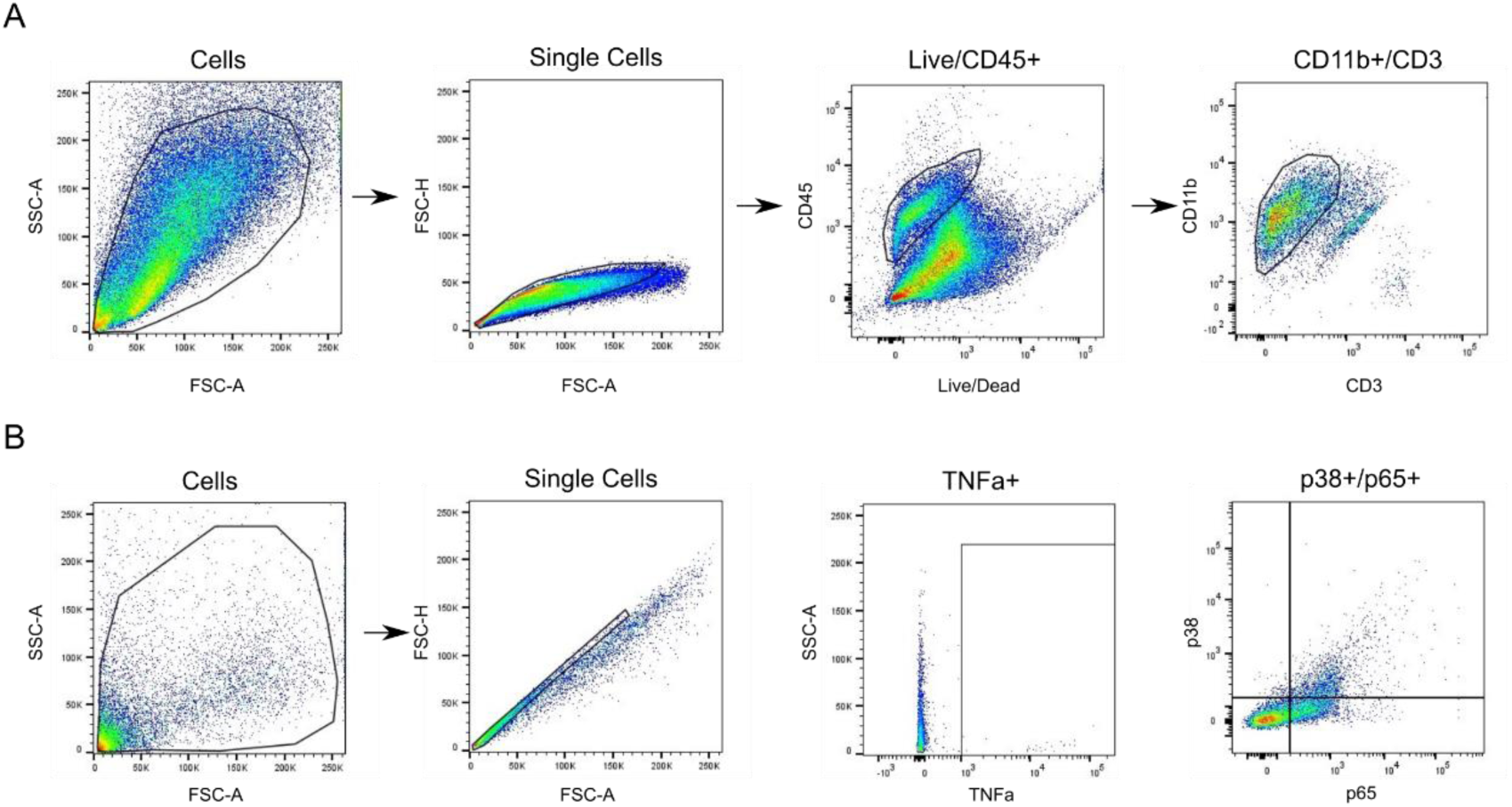
Gating strategies used in flow cytometry experiments. A. Depiction of the gating strategy used to isolate human dorsal root ganglia myeloid cells using fluorescently activated cell sorting. B. Depiction of the gating strategy used to measure TNFα production and phosphorylation of p38 and p65.

**SFigure 10.**
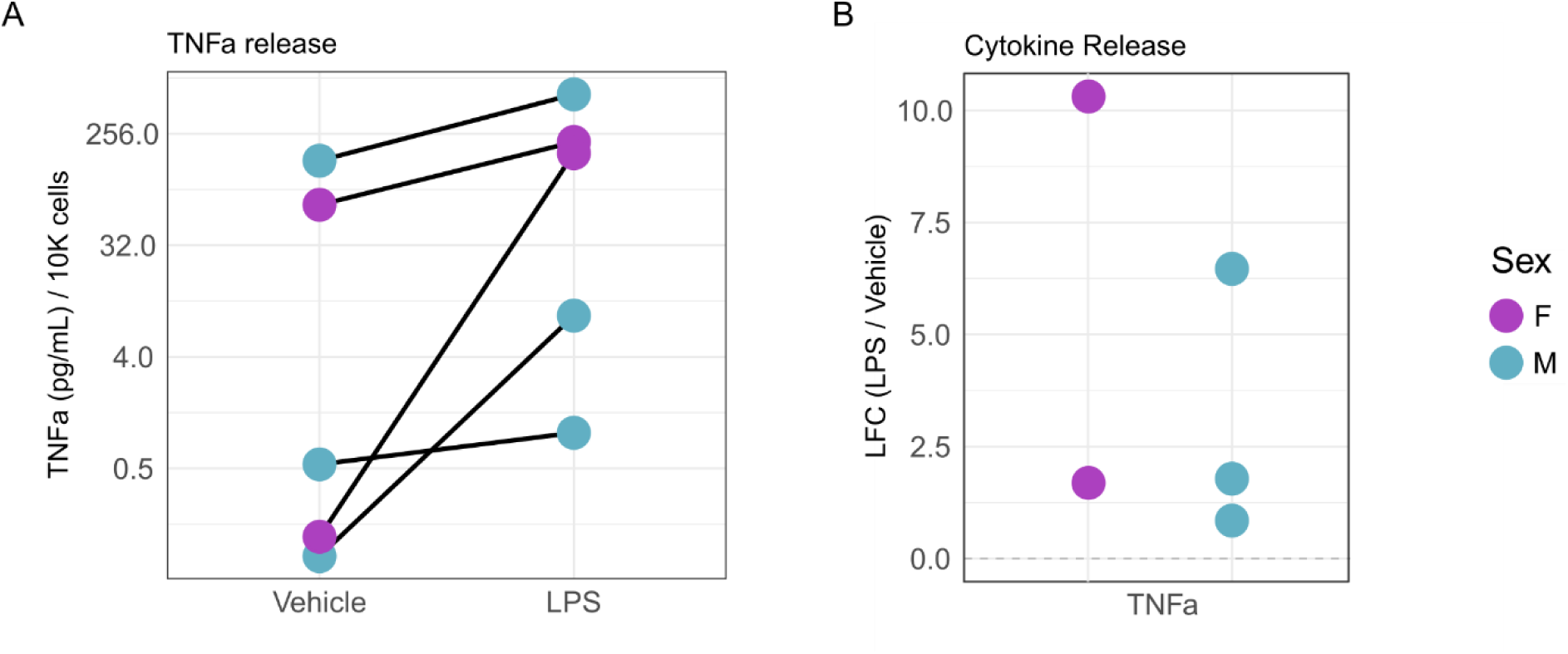
TNFα is released from stimulated myeloid cells from DRG parenchyma. A. TNFα (pg/mL), normalized per 10000 cells plated from Vehicle-treated or LPS-stimulated cultures. B. Log2 fold change (LFC) of TNFα amount from LPS / Vehicle cultures (n = 2F + 3M).

**SFigure 11.**
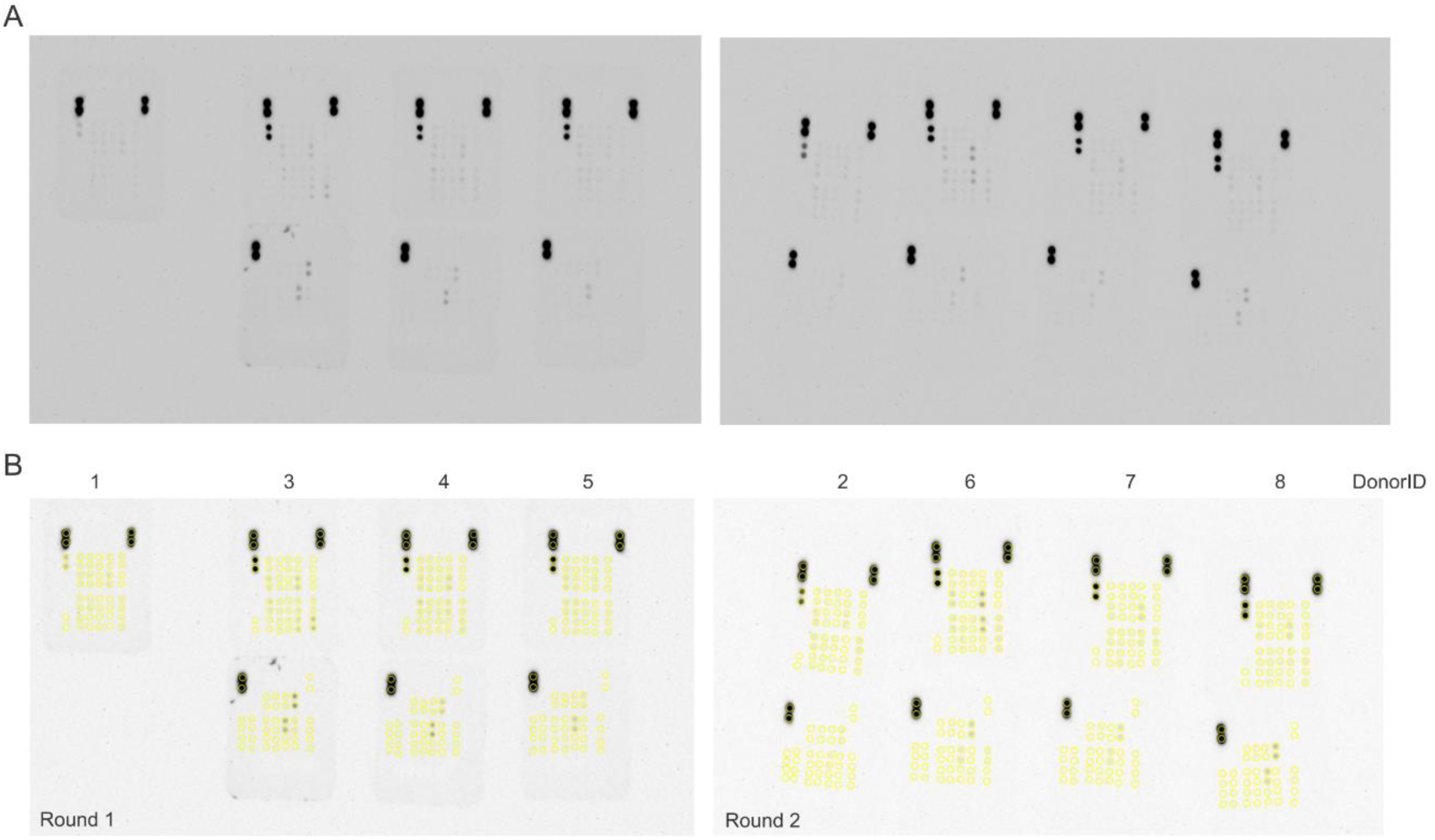
Phosphorylation Array membranes. A. 240 second exposure for round 1 (left) and 2 (right), 2M/2F per round (age-matched, except for Donor 1, where only membrane “A” was processed). B. Selected ROIs (in yellow) for each phosphorylation site, donor numbers match “MS-” donors in Table 1. Contrast enhancement for visualization only. Corresponding source data in Supplemental Table 9.

## Bibliography

[1] Agbanoma G, Li C, Ennis D, Palfreeman AC, Williams LM, Brennan FM. Production of TNF-α in Macrophages Activated by T Cells, Compared with Lipopolysaccharide, Uses Distinct IL-10–Dependent Regulatory Mechanism. J Immunol 2012;188.

[2] Alexander SPH, Mathie AA, Peters JA, Veale EL, Striessnig J, Kelly E, Armstrong JF, Faccenda E, Harding SD, Davies JA, Aldrich RW, Attali B, Baggetta AM, Becirovic E, Biel M, Bill RM, Caceres AI, Catterall WA, Conner AC, Davies P, De Clerq K, Delling M, Di Virgilio F, Falzoni S, Fenske S, Fortuny-Gomez A, Fountain S, George C, Goldstein SAN, Grimm C, Grissmer S, Ha K, Hammelmann V, Hanukoglu I, Hu M, Ijzerman AP, Jabba S V., Jarvis M, Jensen AA, Jordt SE, Kaczmarek LK, Kellenberger S, Kennedy C, King B, Kitchen P, Liu Q, Lynch JW, Meades J, Mehlfeld V, Nicke A, Offermanns S, Perez-Reyes E, Plant LD, Rash L, Ren D, Salman MM, Sieghart W, Sivilotti LG, Smart TG, Snutch TP, Tian J, Trimmer JS, Van den Eynde C, Vriens J, Wei AD, Winn BT, Wulff H, Xu H, Yang F, Fang W, Yue L, Zhang X, Zhu M. The Concise Guide to PHARMACOLOGY 2023/24: Ion channels. Br J Pharmacol 2023;180.

[3] Bair E, Hastie T, Paul D, Tibshirani R. Prediction by Supervised Principal Components. 10.1198/016214505000000628 2006;101:119–137. doi:10.1198/016214505000000628.

[4] Barry A, Schmidt M, Price T. Bulk proteomics of naive human dorsal root ganglion (Version 1). 2025. doi:10.26275/Z7UY-KUIF.

[5] Barry AM, Price TJ, Schmidt M. Bulk Proteomics (DIA-MS) of Human Dorsal Root Ganglion. n.d. doi:10.17504/protocols.io.j8nlk8j56l5r/v1.

[6] Barry AM, Sondermann JR, Sondermann J-H, Gomez-Varela D, Schmidt M. Region-resolved quantitative proteome profiling reveals molecular dynamics associated with chronic pain in the PNS and spinal cord. Front Mol Neurosci 2018;11:259.

[7] Barry AM, Zhao N, Yang X, Bennett DL, Baskozos G. Deep RNA-seq of male and female murine sensory neuron subtypes after nerve injury. Pain 2023;164:2196–2215. doi:10.1101/2022.11.21.516781.

[8] Berkley KJ. Sex differences in pain. Behav Brain Sci 1997;20:371–380. doi:10.1017/S0140525X97221485.

[9] Bhuiyan SA, Xu M, Yang L, Semizoglou E, Bhatia P, Pantaleo KI, Tochitsky I, Jain A, Erdogan B, Blair S, Cat V, Mwirigi JM, Sankaranarayanan I, Tavares-Ferreira D, Green U, McIlvried LA, Copits BA, Bertels Z, Del Rosario JS, Widman AJ, Slivicki RA, Yi J, Sharif-Naeini R, Woolf CJ, Lennerz JK, Whited JL, Price TJ, Gereau RW, Renthal W. Harmonized cross-species cell atlases of trigeminal and dorsal root ganglia. Sci Adv 2024;10.

[10] Van Den Bosch F, Manger B, Goupille P, McHugh N, Rødevand E, Holck P, Van Vollenhoven RF, Leirisalo-Repo M, Fitzgerald O, Kron M, Frank M, Kary S, Kupper H. Effectiveness of adalimumab in treating patients with active psoriatic arthritis and predictors of good clinical responses for arthritis, skin and nail lesions. Ann Rheum Dis 2010;69.

[11] Campbell J, Ciesielski CJ, Hunt AE, Horwood NJ, Beech JT, Hayes LA, Denys A, Feldmann M, Brennan FM, Foxwell BMJ. A Novel Mechanism for TNF-α Regulation by p38 MAPK: Involvement of NF-κB with Implications for Therapy in Rheumatoid Arthritis. J Immunol 2004;173.

[12] Chimenti MS, Triggianese P, Conigliaro P, Tonelli M, Gigliucci G, Novelli L, Teoli M, Perricone R. A 2-year observational study on treatment targets in psoriatic arthritis patients treated with TNF inhibitors. Clin Rheumatol 2017;36.

[13] Cignarella A, Vegeto E, Bolego C, Trabace L, Conti L, Ortona E. Sex-oriented perspectives in immunopharmacology. Pharmacol Res 2023;197.

[14] Crandall W V, Mackner LM. Infusion reactions to infliximab in children and adolescents: frequency, outcome and a predictive model. n.d. doi:10.1046/j.0269-2813.2003.01411.x.

[15] Dawes JM, Bennett DL. Addressing the gender pain gap. Neuron 2021;109:2641–2642.

[16] Demichev V, Messner CB, Vernardis SI, Lilley KS, Ralser M. DIA-NN: neural networks and interference correction enable deep proteome coverage in high throughput. Nat Methods 2020;17.

[17] Demichev V, Szyrwiel L, Yu F, Teo GC, Rosenberger G, Niewienda A, Ludwig D, Decker J, Kaspar-Schoenefeld S, Lilley KS, Mülleder M, Nesvizhskii AI, Ralser M. dia-PASEF data analysis using FragPipe and DIA-NN for deep proteomics of low sample amounts. Nat Commun 2022;13.

[18] Dolgalev I. msigdbr: MSigDB Gene Sets for Multiple Organisms in a Tidy Data Format version 7.4.1 from CRAN. 2021 Available: https://rdrr.io/cran/msigdbr/. Accessed 28 Mar 2022.

[19] Doty M, Yun S, Wang Y, Hu M, Cassidy M, Hall B, Kulkarni AB. Integrative multiomic analyses of dorsal root ganglia in diabetic neuropathic pain using proteomics, phospho-proteomics, and metabolomics. Sci Rep 2022;12.

[20] Durinck S, Moreau Y, Kasprzyk A, Davis S, De Moor B, Brazma A, Huber W. BioMart and Bioconductor: A powerful link between biological databases and microarray data analysis. Bioinformatics 2005;21.

[21] Dyring-Andersen B, Løvendorf MB, Coscia F, Santos A, Møller LBP, Colaço AR, Niu L, Bzorek M, Doll S, Andersen JL, Clark RA, Skov L, Teunissen MBM, Mann M. Spatially and cell-type resolved quantitative proteomic atlas of healthy human skin. Nat Commun 2020;11.

[22] Eid SA, Rumora AE, Beirowski B, Bennett DL, Hur J, Savelieff MG, Feldman EL. New perspectives in diabetic neuropathy. Neuron 2023;111.

[23] Farah A, Patel R, Poplawski P, Wastie BJ, Tseng M, Barry AM, Daifallah O, Dubb A, Paul I, Lao Cheng H, Feroz F, Su Y, Chan M, Zeilhofer HU, Price T, Bennett DL, Bannister K, Dawes JM. A role for leucine-rich, glioma inactivated 1 in regulating pain sensitivity. Brain 2024. doi:10.1093/brain/awae302/7762306.

[24] Fawzi MS, El-Shal AS, Rashad NM, Fathy HA. Influence of tumor necrosis factor alpha gene promoter polymorphisms and its serum level on migraine susceptibility in Egyptian patients. J Neurol Sci 2015;348.

[25] Franco-Enzástiga Ú, Inturi NN, Natarajan K, Mwirigi JM, Mazhar K, Schlachetzki JCM, Schumacher M, Price TJ. Epigenomic landscape of the human dorsal root ganglion: sex differences and transcriptional regulation of nociceptive genes Running title: epigenome of the human DRG. Pain 2024. doi:10.1101/2024.03.27.587047.

[26] Furman D, Hejblum BP, Simon N, Jojic V, Dekker CL, Thiebaut R, Tibshirani RJ, Davis MM. Systems analysis of sex differences reveals an immunosuppressive role for testosterone in the response to influenza vaccination. Proc Natl Acad Sci U S A 2014;111.

[27] Gillet LC, Navarro P, Tate S, Röst H, Selevsek N, Reiter L, Bonner R, Aebersold R. Targeted data extraction of the MS/MS spectra generated by data-independent acquisition: A new concept for consistent and accurate proteome analysis. Mol Cell Proteomics 2012;11.

[28] Greenspan JD, Craft RM, LeResche L, Arendt-Nielsen L, Berkley KJ, Fillingim RB, Gold MS, Holdcroft A, Lautenbacher S, Mayer EA, Mogil JS, Murphy AZ, Traub RJ. Studying sex and gender differences in pain and analgesia: A consensus report. Pain 2007;132.

[29] Hadizadeh F, Johansson T, Johansson A, Karlsson T, Ek WE. Effects of oral contraceptives and menopausal hormone therapy on the risk of rheumatoid arthritis: a prospective cohort study. Rheumatol (United Kingdom) 2024;63.

[30] Huynh L, Kusnadi A, Park SH, Murata K, Park-Min KH, Ivashkiv LB. Opposing regulation of the late phase TNF response by mTORC1-IL-10 signaling and hypoxia in human macrophages. Sci Rep 2016;6.

[31] Hyrich KL, Watson KD, Silman AJ, Symmons DPM. Predictors of response to anti-TNF-α therapy among patients with rheumatoid arthritis: Results from the British Society for Rheumatology Biologics Register. Rheumatology 2006;45.

[32] Ikegami D, Navratilova E, Yue X, Moutal A, Kopruszinski CM, Khanna R, Patwardhan A, Dodick DW, Porreca F. A prolactin-dependent sexually dimorphic mechanism of migraine chronification. Cephalalgia 2022;42:197–208. doi:10.1177/03331024211039813.

[33] Jain A, Gyori BM, Hakim S, Jain A, Sun L, Petrova V, Bhuiyan SA, Zhen S, Wang Q, Kawaguchi R, Bunga S, Taub DG, Ruiz-Cantero MC, Tong-Li C, Andrews N, Kotoda M, Renthal W, Sorger PK, Woolf CJ. Nociceptor-immune interactomes reveal insult-specific immune signatures of pain. Nat Immunol 2024.

[34] Jung M, Dourado M, Maksymetz J, Jacobson A, Laufer BI, Baca M, Foreman O, Hackos DH, Riol-Blanco L, Kaminker JS. Cross-species transcriptomic atlas of dorsal root ganglia reveals species-specific programs for sensory function. Nat Commun 2023;14.

[35] Kamada M, Irahara M, Maegawa M, Yasui T, Takeji T, Yamada M, Tezuka M, Kasai Y, Aono T. Effect of hormone replacement therapy on post-menopausal changes of lymphocytes and T cell subsets. J Endocrinol Invest 2000;23.

[36] LaCroix-Fralish ML, Ledoux JB, Mogil JS. The Pain Genes Database: An interactive web browser of pain-related transgenic knockout studies. Pain 2007;131.

[37] Lakshmikanth T, Consiglio C, Sardh F, Forlin R, Wang J, Tan Z, Barcenilla H, Rodriguez L, Sugrue J, Noori P, Páez LP, Gonzalez L, Mugabo CH, Johnsson A, Hallgren Å, Pou C, Chen Y, Mikeš J, James A, Dahlqvist P, Wahlberg J, Hagelin A, Holmberg M, Degerblad M, Isaksson M, Duffy D, Kämpe O, Landegren N, Brodin P. Immune system adaptation during gender-affirming testosterone treatment. Nature 2024;159:155–164.

[38] Love MI, Huber W, Anders S. Moderated estimation of fold change and dispersion for RNA-seq data with DESeq2. Genome Biol 2014 1512 2014;15:1–21. doi:10.1186/S13059-014-0550-8.

[39] Maguire AD, Friedman TN, Villarreal Andrade DN, Haq F, Dunn J, Pfeifle K, Tenorio G, Buro K, Plemel JR, Kerr BJ. Sex differences in the inflammatory response of the mouse DRG and its connection to pain in experimental autoimmune encephalomyelitis. Sci Rep 2022;12.

[40] Meloto CB, Benavides R, Lichtenwalter RN, Wen X, Tugarinov N, Zorina-Lichtenwalter K, Chabot-Doré AJ, Piltonen MH, Cattaneo S, Verma V, Klares R, Khoury S, Parisien M, Diatchenko L. Human pain genetics database: A resource dedicated to human pain genetics research. Pain 2018;159.

[41] Mogil JS. Sex differences in pain and pain inhibition: multiple explanations of a controversial phenomenon. 2012 doi:10.1038/nrn3360.

[42] Ochoa D, Hercules A, Carmona M, Suveges D, Baker J, Malangone C, Lopez I, Miranda A, Cruz-Castillo C, Fumis L, Bernal-Llinares M, Tsukanov K, Cornu H, Tsirigos K, Razuvayevskaya O, Buniello A, Schwartzentruber J, Karim M, Ariano B, Martinez Osorio RE, Ferrer J, Ge X, Machlitt-Northen S, Gonzalez-Uriarte A, Saha S, Tirunagari S, Mehta C, Roldán-Romero JM, Horswell S, Young S, Ghoussaini M, Hulcoop DG, Dunham I, Mcdonagh EM. The next-generation Open Targets Platform: reimagined, redesigned, rebuilt. Nucleic Acids Res 2023;51.

[43] Perez-Riverol Y, Bai J, Bandla C, García-Seisdedos D, Hewapathirana S, Kamatchinathan S, Kundu DJ, Prakash A, Frericks-Zipper A, Eisenacher M, Walzer M, Wang S, Brazma A, Vizcaíno JA. The PRIDE database resources in 2022: A hub for mass spectrometry-based proteomics evidences. Nucleic Acids Res 2022;50.

[44] Rabilloud T. Membrane proteins and proteomics: Love is possible, but so difficult. Electrophoresis 2009;30.

[45] Ray PR, Shiers S, Caruso JP, Tavares-Ferreira D, Sankaranarayanan I, Uhelski ML, Li Y, North RY, Tatsui C, Dussor G, Burton MD, Dougherty PM, Price TJ. RNA profiling of human dorsal root ganglia reveals sex differences in mechanisms promoting neuropathic pain. Brain 2023.

[46] Ritchie ME, Phipson B, Wu D, Hu Y, Law CW, Shi W, Smyth GK. Limma powers differential expression analyses for RNA-sequencing and microarray studies. Nucleic Acids Res 2015;43.

[47] Rusman T, van Vollenhoven RF, van der Horst-Bruinsma IE. Gender Differences in Axial Spondyloarthritis: Women Are Not So Lucky. Curr Rheumatol Rep 2018;20.

[48] Salzer J, Feltri ML, Jacob C. Schwann Cell Development and Myelination. Cold Spring Harb Perspect Biol 2024;16.

[49] Schultheiss JPD, Brand EC, Lamers E, van den Berg WCM, van Schaik FDM, Oldenburg B, Fidder HH. Earlier discontinuation of TNF-α inhibitor therapy in female patients with inflammatory bowel disease is related to a greater risk of side effects. Aliment Pharmacol Ther 2019;50.

[50] Schwaid AG, Krasowka-Zoladek A, Chi A, Cornella-Taracido I. Comparison of the Rat and Human Dorsal Root Ganglion Proteome. Sci Rep 2018;8:1–10.

[51] Sorge RE, LaCroix-Fralish ML, Tuttle AH, Sotocinal SG, Austin JS, Ritchie J, Chanda ML, Graham AC, Topham L, Beggs S, Salter MW, Mogil JS. Spinal cord toll-like receptor 4 mediates inflammatory and neuropathic hypersensitivity in male but not female mice. J Neurosci 2011;31:15450–15454.

[52] Sorge RE, Mapplebeck JCS, Rosen S, Beggs S, Taves S, Alexander JK, Martin LJ, Austin JS, Sotocinal SG, Chen D, Yang M, Shi XQ, Huang H, Pillon NJ, Bilan PJ, Tu Y, Klip A, Ji RR, Zhang J, Salter MW, Mogil JS. Different immune cells mediate mechanical pain hypersensitivity in male and female mice. Nat Neurosci 2015;18.

[53] Souto A, Maneiro JR, Gómez-Reino JJ. Rate of discontinuation and drug survival of biologic therapies in rheumatoid arthritis: A systematic review and meta-analysis of drug registries and health care databases. Rheumatol (United Kingdom) 2016;55.

[54] Suzuki M, Tetsuka T, Yoshida S, Watanabe N, Kobayashi M, Matsui N, Okamoto T. The role of p38 mitogen-activated protein kinase in IL-6 and IL-8 production from the TNF-α-or IL-1β-stimulated rheumatoid synovial fibroblasts. FEBS Lett 2000;465.

[55] Tavares-Ferreira D, Shiers S, Ray PR, Wangzhou A, Jeevakumar V, Sankaranarayanan I, Cervantes AM, Reese JC, Chamessian A, Copits BA, Dougherty PM, Gereau RW, Burton MD, Dussor G, Price TJ. Spatial transcriptomics of dorsal root ganglia identifies molecular signatures of human nociceptors. Sci Transl Med 2022;14:eabj8186. doi:DOI: 10.1126/scitranslmed.abj8186.

[56] Themistocleous AC, Baskozos G, Blesneac I, Comini M, Megy K, Chong S, Deevi SV V, Ginsberg L, Gosal D, Hadden RDM, Horvath R, Mahdi-Rogers M, Manzur A, Mapeta R, Marshall A, Matthews E, McCarthy MI, Reilly MM, Renton T, Rice ASC, Vale TA, van Zuydam N, Walker SM, Woods CG, Bennett DLH. Investigating genotype–phenotype relationship of extreme neuropathic pain disorders in a UK national cohort. Brain Commun 2023;5. doi:10.1093/braincomms/fcad037.

[57] De Vito R, Bellio R, Trippa L, Parmigiani G. Multi-study Factor Analysis. 2016. Available: http://arxiv.org/abs/1611.06350.

[58] Wangzhou A, Paige C, Neerukonda S V., Naik DK, Kume M, David ET, Dussor G, Ray PR, Price TJ. A ligand-receptor interactome platform for discovery of pain mechanisms and therapeutic targets. Sci Signal 2021;14.

[59] de Winter JCF, Dodou D, Wieringa PA. Exploratory factor analysis with small sample sizes. Multivariate Behav Res 2009;44:147–181.

[60] Wu CC, Yates JR. The application of mass spectrometry to membrane proteomics. Nat Biotechnol 2003;21.

[61] Wu T, Hu E, Xu S, Chen M, Guo P, Dai Z, Feng T, Zhou L, Tang W, Zhan L, Fu X, Liu S, Bo X, Yu G. clusterProfiler 4.0: A universal enrichment tool for interpreting omics data. Innov 2021;2:100141.

[62] Yu X, Liu H, Hamel KA, Morvan MG, Yu S, Leff J, Guan Z, Braz JM, Basbaum AI. Dorsal root ganglion macrophages contribute to both the initiation and persistence of neuropathic pain. Nat Commun 2020;11:1–12.

[63] Zelinkova Z, Bultman E, Vogelaar L, Bouziane C, Kuipers EJ, van der Woude CJ. Sex-dimorphic adverse drug reactions to immune suppressive agents in inflammatory bowel disease. World J Gastroenterol 2012;18.

[64] Zhao N, Bennett DL, Baskozos G, Barry AM. Predicting pain genes: multi-modal data integration using probabilistic classifiers and interaction networks. Bioinforma Adv 2024;4:0–0.

[65] Zheng Y, Liu P, Bai L, Trimmer JS, Bean BP, Ginty DD. Deep Sequencing of Somatosensory Neurons Reveals Molecular Determinants of Intrinsic Physiological Properties. Neuron 2019;103:598–616.e7. doi:10.1016/j.neuron.2019.05.039.

[66] Zhu A, Ibrahim JG, Love MI. Heavy-tailed prior distributions for sequence count data: removing the noise and preserving large differences. Bioinformatics 2019;35:2084–2092. doi:10.1093/BIOINFORMATICS/BTY895.

